# zsasa: a Zig-based engine for high-throughput solvent accessible surface area at proteome scale

**DOI:** 10.64898/2026.06.29.733683

**Authors:** Tsubasa Nagae, Kentaro Tomii

## Abstract

Advances in structural and computational biology now yield structures by the hundreds of thousands, from predicted-structure databases to molecular-dynamics trajectories. Solvent accessible surface area (SASA), a long-established descriptor of protein stability, binding, and interfaces, remains central to their analysis. Established SASA tools are still widely used, but each covers only part of what such data now demand in input formats, outputs, and speed. We present `zsasa`, a fast, full-featured, open-source SASA engine for single structures, proteome-scale collections, and trajectories, usable from the command line or Python. It uses the same algorithms as established tools and owes its speed to the implementation, which makes full use of multicore processors and SIMD instructions while keeping memory use low. On a single laptop, `zsasa` reproduced FreeSASA, a widely used SASA library, down to individual atom areas on 4,370 *Escherichia coli* AlphaFold models. On the 23,586 structures of the human AlphaFold proteome, it was 3.3-fold faster than FreeSASA run in parallel while using a third or less of the peak memory of the other tools tested, and an optional approximate mode was 10-fold faster, with total SASA within 0.7% of the default results on average. On molecular-dynamics trajectories, it was 12-to 52-fold faster than MDTraj. As a demonstration, we computed interface burial for more than 600,000 predicted human protein dimers in under half an hour and found that a large buried area did not imply a confident predicted interface. `zsasa` is available under the MIT license at https://github.com/N283T/zsasa.

**Impact statement:** Solvent accessible surface area is among the most widely used structural descriptors, but established calculators each cover only part of what proteome-scale collections of predicted structures and long trajectories now demand. `zsasa` reproduces FreeSASA Shrake–Rupley results, processes tens of thousands of structures per minute on a laptop using little memory, and writes tables that can be joined directly with other annotations.

## Introduction

Solvent accessible surface area (SASA) measures the portion of a molecular surface that is reachable by a solvent probe. Although simple to define, SASA recurs throughout structural biology: residue exposure, protein stability, ligand binding, mutation effects, conformational change, and protein–protein interfaces all depend on surface accessibility in some form. Buried surface area extends the same idea to complexes by comparing isolated partners with the assembled state. Because SASA depends on probe radius, atomic radii, hydrogen handling, and sampling density, reproducible calculation requires well-defined algorithms and recorded settings. The Shrake–Rupley and Lee–Richards algorithms remain standard choices because they are interpretable, parameterizable, and straightforward to compare across implementations (Shrake and Rupley 1973; Lee and Richards 1971). Analytical and molecular-surface formulations clarify the distinctions among accessible surfaces, excluded surfaces, and exact analytical alternatives (Connolly 1983; Richmond 1984; Fraczkiewicz and Braun 1998).

The scale of structural data has changed. A single project may now involve directories of experimental structures, cryo-electron-microscopy assemblies, predicted models, designed proteins, and molecular-dynamics trajectories. Proteome-scale predicted-structure collections such as the AlphaFold Database (AFDB) have made directory-level analysis of tens of thousands of models a routine starting point (Jumper et al. 2021; Varadi et al. 2022, 2024; Bertoni et al. 2026), and predicted complex datasets extend the same scaling pressure to multimeric structures (Schweke et al. 2024; Han et al. 2026; Qi et al. 2026). In these settings, SASA is a feature that must be generated reproducibly across large collections for downstream statistical, machine-learning, or triage analyses, as in de novo binder prioritization, where interface descriptors such as ΔSASA and shape complementarity complement structure-prediction confidence scores (Overath et al. 2025). At this scale, the surface-area kernel is only part of the cost: files in several formats must be parsed, settings applied consistently, memory kept bounded for very large structures, and outputs written in a form that joins with residue metadata. Reproducibility, the ability to record how a number was produced and to regenerate it later, then depends on the tool as much as on the user’s notes.

Existing tools address parts of this problem (Table 1). FreeSASA provides a mature, widely used open-source reference implementation (Mitternacht 2016), and more recent projects such as RustSASA and Lahuta have explored SIMD acceleration and bitmask lookup-table optimization (Campbell 2026; Sejdiu and Babu 2026). Earlier approximate atomic-surface methods also made the tradeoff between speed and agreement a recurring design theme in SASA software (Weiser, Shenkin, and Still 1999; Durham et al. 2009). These tools remain valuable, but large and heterogeneous workloads call for an implementation designed from the start for high-throughput execution, with explicit workflow files, consistent output schemas, selectable precision and threading, and atom-radius assignment that remains practical for ligands, cofactors, and modified components.

**Table 1.**
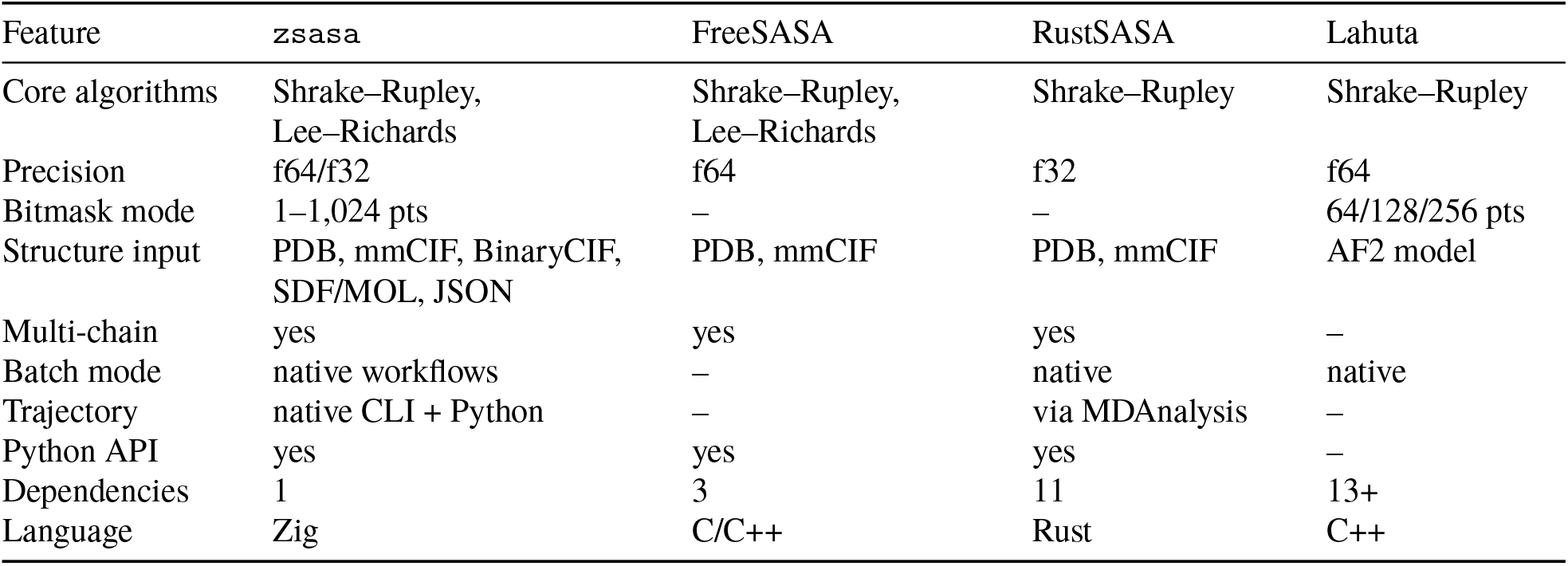
SASA capabilities of the compared tools. Entries summarize SASA-relevant functionality checked against the source code of each tool and the `zsasa` documentation. For Lahuta, entries refer to its `sasa-sr` mode, which requires AlphaFold2 (AF2) models as input, processes only chain A, and supports bitmask calculation at 64, 128, or 256 points. FreeSASA has no native directory/batch mode. RustSASA Python and MDAnalysis support are provided through companion packages; dependency counts are explained in Supplementary Note S1. “–” indicates that the feature is not available.

We developed `zsasa` for this setting and evaluate it on three criteria. First, *numerical agreement*: under matched settings, the exact mode should reproduce a reference implementation, and the optional approximate mode should have a measured error. Second, *throughput and memory*: the engine should process proteome-scale directories and trajectories quickly and with low peak memory, and very large single structures should remain tractable. Third, *integration*: compressed-input handling, workflow files, configurable classification, and structured outputs should let SASA be computed as one step of a larger analysis. `zsasa` introduces no new definition of surface area: it implements the standard algorithms reproducibly and owes its speed to parallel execution, vectorization, and memory handling. We evaluate it on static structures, molecular-dynamics trajectories, and two sets of predicted human dimer models that together contain more than 600,000 structures.

## Results

### Design of zsasa

`zsasa` is a SASA calculation engine written primarily in Zig (https://ziglang.org/) and available through command-line, C ABI, and Python interfaces. Figure 1 illustrates the quantity being computed and summarizes the supported workloads, interfaces, and outputs.

**Figure 1.**
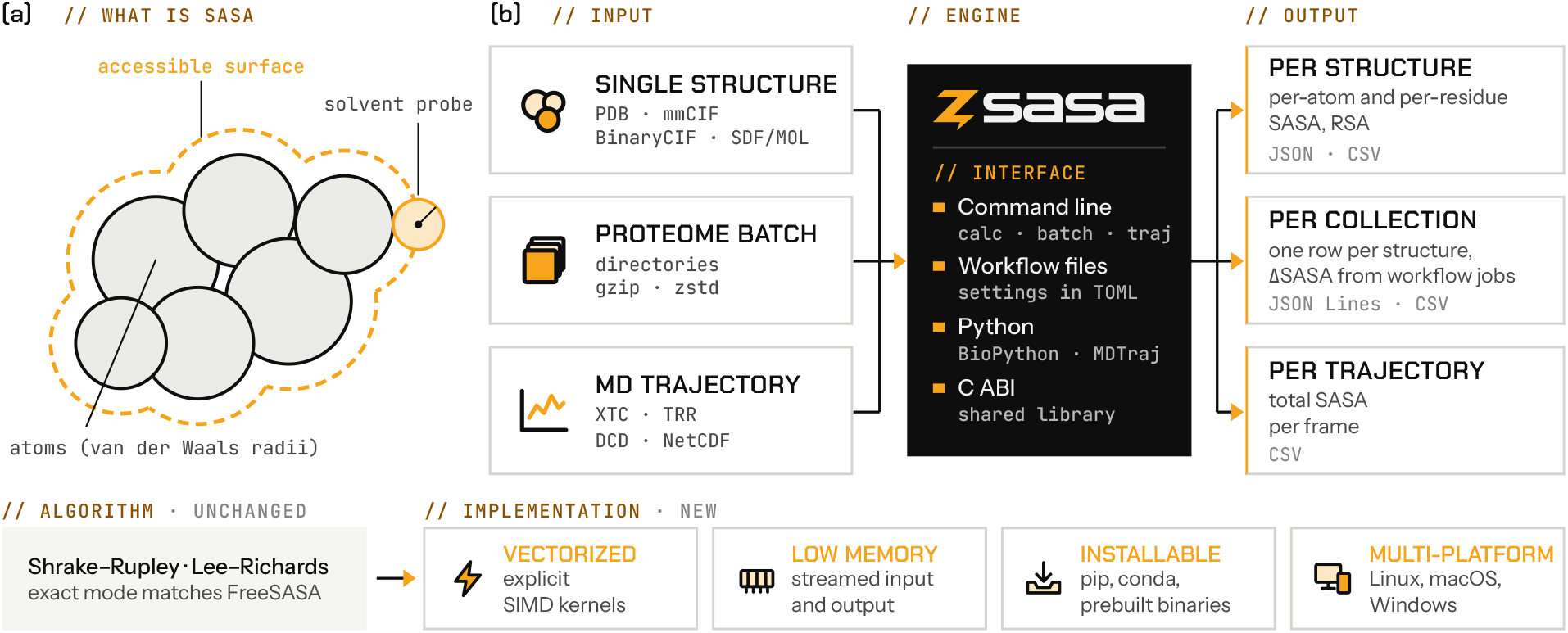
Overview of zsasa. (a) Schematic of the solvent accessible surface, traced by the center of a solvent probe rolled over the van der Waals surface of the atoms. (b) One engine, driven from the command line, workflow files that record the settings of a run, the Python API, or the C ABI, serves single structures, proteome-scale directories, and molecular-dynamics trajectories, and writes per-structure, per-collection, and per-frame results. Bottom: the algorithms are those of existing tools, and the exact mode reproduces FreeSASA (Figure 2a); the implementation is new.

In the default exact mode, `zsasa` computes Shrake–Rupley SASA by sampling points on solvent-expanded atomic spheres, with neighbor lists to avoid unnecessary atom-pair checks (Shrake and Rupley 1973); the Lee–Richards algorithm is available as an independent slice-integration path (Lee and Richards 1971). For throughput-oriented workloads, an optional Shrake–Rupley bitmask mode precomputes per-point occlusion masks and combines them with bitwise operations. The lookup-table design was inspired by Lahuta (Sejdiu and Babu 2026) but is implemented independently and accepts any point count from 1 to 1,024. Precision is selectable between double (f64) and single (f32).

Because the exact-mode calculation is the same as in existing Shrake–Rupley implementations, the speedups reported below come from implementation choices such as data layout, allocation behavior, vectorization, and parallel scheduling. Supplementary Note S1 discusses what Zig contributes to them.

Atomic radii are assigned by built-in classifiers or by custom tables; the default classifier extends ProtOr-style radii to ligands, cofactors, and modified residues using Chemical Component Dictionary (CCD) topology (John D. Westbrook et al. 2015; Tsai et al. 1999). The command-line interface (CLI) reads the common structure formats, including compressed files, and the Python package works with the common structure and trajectory libraries. Settings can be recorded in TOML workflow files that define named jobs, such as chain A, chain B, and complex AB for buried-surface analyses, and batch results are streamed as JSON Lines with optional residue-level maps. Table 1 compares these capabilities with those of the other tools; implementation details are given in Materials and Methods and in Supplementary Note S1.

### Benchmark overview

We evaluated `zsasa` v0.12.1 with five benchmark suites (Table 2): validation against FreeSASA, batch throughput, and single-file timing for static structures, and validation against MDTraj and throughput for trajectories. All head-to-head benchmarks were run on one consumer laptop (Apple M4, 4 performance and 6 efficiency cores, 32 GB of memory) with pinned comparator builds (Materials and Methods). Throughout, f64 and f32 denote the exact mode at double and single precision; a result refers to this mode unless it is labeled *bitmask*, which denotes the approximate lookup-table mode. Representative results are summarized in Table 3.

**Table 2.**
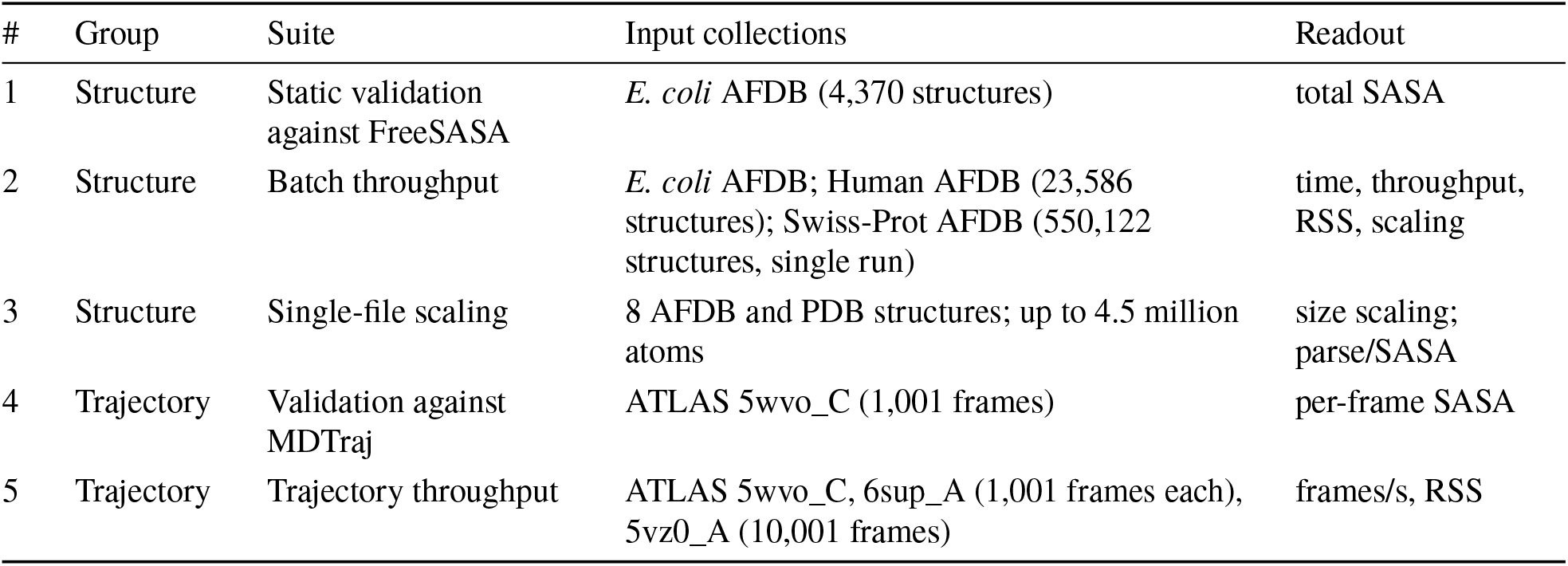
Benchmark suites and input collections. Detailed settings and measurement fields are provided in the Supplement. AFDB = AlphaFold Database; RSS = resident set size.

**Table 3.**
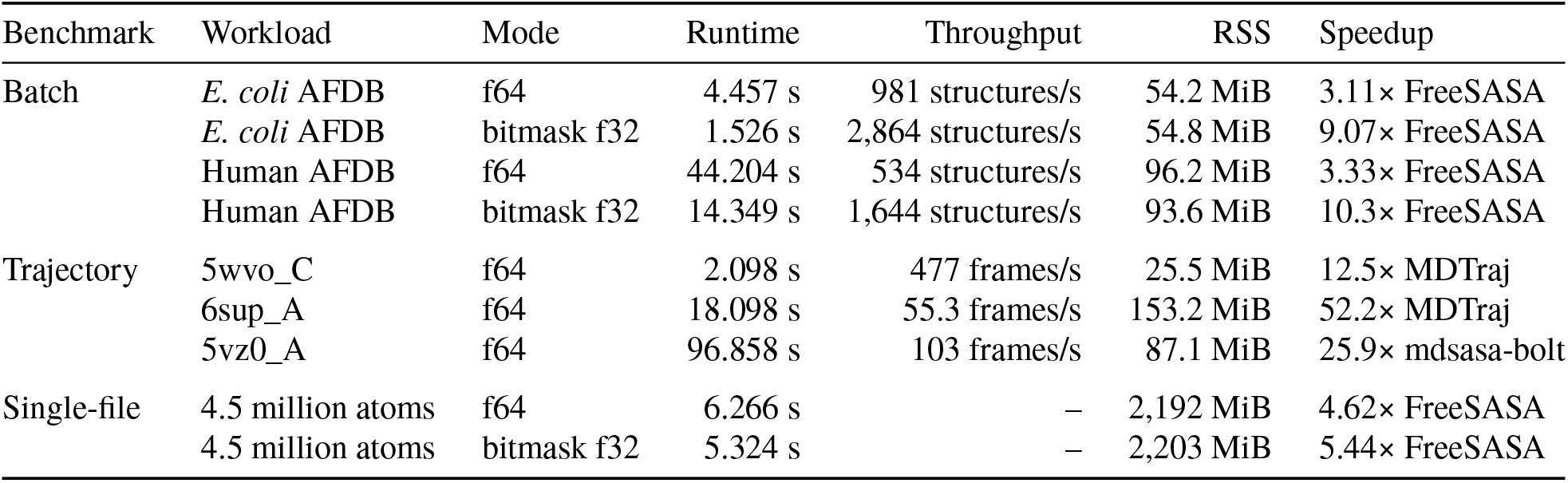
Representative performance results. (`zsasa` v0.12.1, 10 threads). Batch and trajectory runs used 128 sphere points and single-file runs used 100. RSS values are peak RSS; speedups are relative to the named comparator.

| Benchmark | Workload | Mode | Runtime | Throughput | RSS | Speedup |
|---|---|---|---|---|---|---|
| Batch | *E. coli* AFDB | f64 | 4.457 s | 981 structures/s | 54.2 MiB | 3.11× FreeSASA |
|  | *E. coli* AFDB | bitmask f32 | 1.526 s | 2,864 structures/s | 54.8 MiB | 9.07× FreeSASA |
|  | Human AFDB | f64 | 44.204 s | 534 structures/s | 96.2 MiB | 3.33× FreeSASA |
|  | Human AFDB | bitmask f32 | 14.349 s | 1,644 structures/s | 93.6 MiB | 10.3× FreeSASA |
| Trajectory | 5wvo_C | f64 | 2.098 s | 477 frames/s | 25.5 MiB | 12.5× MDTraj |
|  | 6sup_A | f64 | 18.098 s | 55.3 frames/s | 153.2 MiB | 52.2× MDTraj |
|  | 5vz0_A | f64 | 96.858 s | 103 frames/s | 87.1 MiB | 25.9× mdsasa-bolt |
| Single-file | 4.5 million atoms | f64 | 6.266 s | – | 2,192 MiB | 4.62× FreeSASA |
|  | 4.5 million atoms | bitmask f32 | 5.324 s | – | 2,203 MiB | 5.44× FreeSASA |

### Structure benchmarks

#### Validation against FreeSASA

Across 4,370 *E. coli* AlphaFold Database structures, the total SASA from the f64 mode agreed with that from FreeSASA at 128 sphere points, with residual *R*^2^ (Table S1) = 1.000000, mean relative difference = 0.000021%, and maximum relative difference = 0.00023% (Figure 2a). The agreement extended beyond the structure totals: all 10,520,167 per-atom areas and 1,349,634 per-residue areas matched FreeSASA at the two-decimal precision of its text output. The f32 mode agreed similarly closely (mean relative difference = 0.00010%, maximum relative difference = 0.015%), and the agreement did not depend on the sphere-point count from 64 to 1,024 (Figure 2b). The agreement is this close because `zsasa` places its sphere points on the same golden-section spiral as FreeSASA. Therefore, the f64 mode is suitable when results must match existing FreeSASA-based analyses. RustSASA, whose golden-section spiral is shifted by half a step along the polar axis and wound in the opposite direction, scattered symmetrically around FreeSASA and converged as the point count increased (mean relative difference 0.28% at 128 points and 0.055% at 1,024 points).

**Figure 2.**
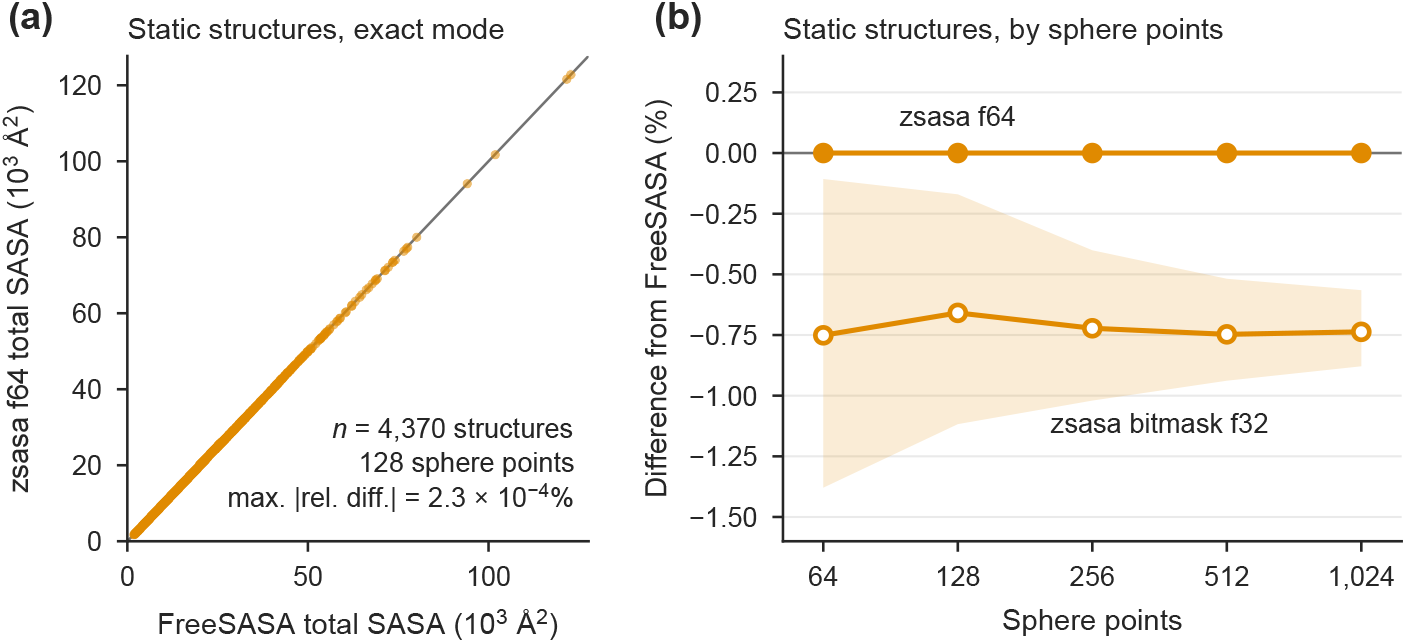
Numerical agreement with FreeSASA. (a) Per-structure total SASA from `zsasa` f64 versus FreeSASA at 128 sphere points for 4,370 *E. coli* AlphaFold Database structures, with the identity line. (b) Signed relative difference from FreeSASA across sphere-point counts for the same structures. Lines show the median and shaded bands span the 5th–95th percentiles; negative values indicate lower total SASA than FreeSASA. Filled markers denote the exact f64 mode and open markers the bitmask f32 mode.

The bitmask mode is a throughput-oriented path with an explicit error envelope (Figure 2b). At 128 sphere points, the bitmask f32 mode retained close agreement with FreeSASA (residual *R*^2^ = 0.999811, mean relative difference = 0.662%, maximum relative difference = 2.02%), and the bitmask f64 mode was virtually identical. The difference is a systematic underestimate rather than random scatter: the median signed relative difference was −0.66% at 128 points (5th–95th percentile −1.12% to −0.17%) and −0.74% at 1,024 points, and Lahuta’s bitmask mode showed the same offset over its supported 64–256 points. The lookup table quantizes directions and angles in fixed steps, which introduces an offset that is not removed by using more sphere points, although the spread across structures narrowed (maximum relative difference 1.08% at 1,024 points). For comparison, the choice of sampling changes exact results by a similar amount: exact `zsasa` at 128 points differed from exact `zsasa` at 1,024 points by 0.20% on average (maximum 1.23%), and RustSASA differed from FreeSASA at 128 points by 0.28% on average (maximum 2.15%). The bitmask offset is about three times the sampling error of a typical 128-point calculation, and its worst case is similar to the difference between two exact implementations. We consider this level suitable for screening, ranking, and feature generation, where SASA serves as a relative descriptor. The full point-count sweep and the secondary 20-slice Lee–Richards check are given in Tables S2–S4 and Figures S1 and S2.

#### Proteome-scale batch throughput

On the 4,370-structure *E. coli* collection at 128 sphere points and 10 threads, `zsasa` f64 ran in 4.457 s (981 structures/s) and bitmask f32 in 1.526 s (2,864 structures/s), corresponding to 3.11-fold and 9.07-fold speedups over the FreeSASA batch wrapper. Peak resident set size (RSS) was 54.2 and 54.8 MiB, respectively. The f64 mode was 1.38-fold faster than RustSASA and 1.58-fold faster than Lahuta’s exact mode, and the bitmask f32 mode was 1.36-fold faster than Lahuta’s bitmask mode. Thread scaling was near-linear at low thread counts and gradually flattened (Figure 3a); the f64 mode reached a 6.60-fold speedup at 10 threads relative to single-threaded execution (parallel efficiency 0.66), and bitmask f32 a 6.70-fold speedup, whereas the FreeSASA wrapper did not scale beyond 4 threads. Parallel efficiency was 0.95 at 4 threads and fell once the 4 performance cores of the test machine were exceeded, so part of the gap from ideal scaling reflects the slower efficiency cores. zsasa is the only tool in the high-throughput, low-memory region of the throughput-versus-memory plot (Figure 3b): the fastest comparator, Lahuta’s bitmask mode, reached 2,110 structures/s at 177 MiB peak RSS, and all comparators used 3.1–4.2 times the peak RSS of `zsasa`. Raw readouts are provided in Tables S5 and S6.

**Figure 3.**
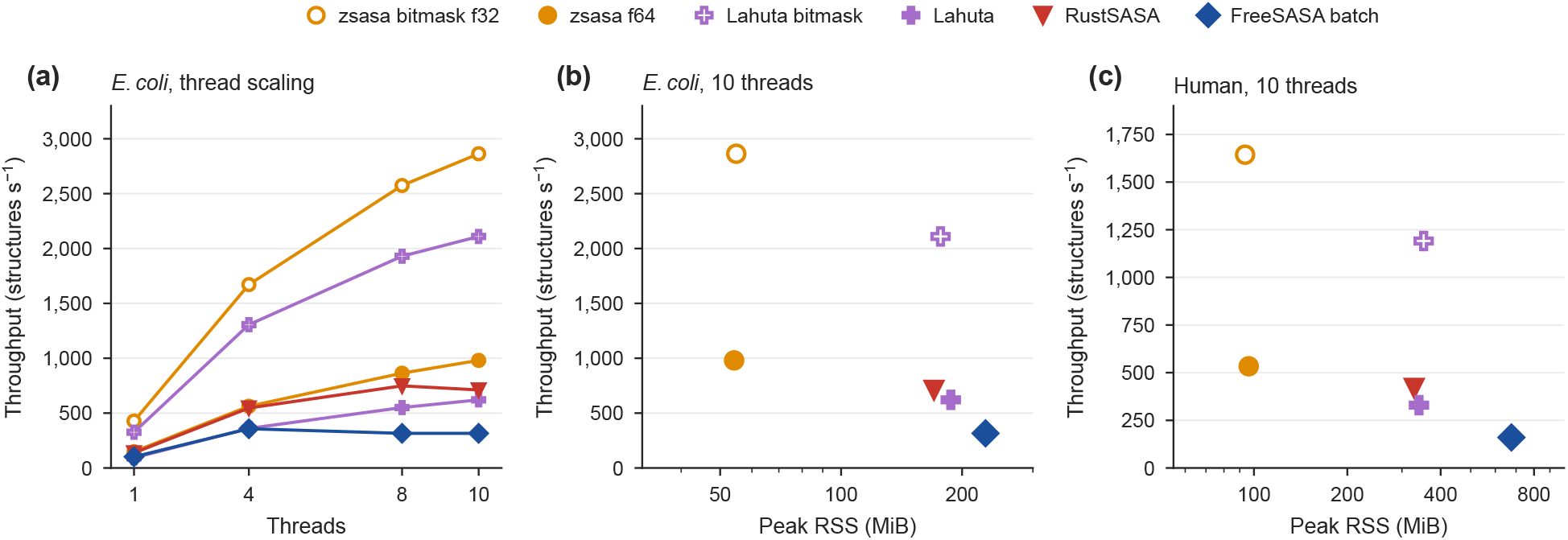
Batch throughput and peak memory on AlphaFold Database collections (128 sphere points). (a) Throughput versus thread count on the *E. coli* collection (4,370 structures). (b, c) Throughput versus peak RSS at 10 threads for (b) the *E. coli* collection and (c) the Human collection (23,586 structures); higher and farther left is better. Values are means of three runs. Filled markers denote exact modes and open markers bitmask modes. FreeSASA batch refers to the multi-file wrapper described in Materials and Methods. `zsasa` f32 and bitmask f64 nearly coincide with f64 and bitmask f32, respectively, and are omitted.

The Human collection contains 23,586 structures, 5.4 times as many as the *E. coli* collection, with larger proteins on average. Throughput was correspondingly lower (f64 44.204 s, 534 structures/s; bitmask f32 14.349 s, 1,644 structures/s; Figure 3c), but the advantage over the comparators held: a 3.33-fold speedup over the FreeSASA wrapper in f64 and a 10.3-fold speedup in bitmask f32, with bitmask f32 1.38-fold faster than Lahuta’s bitmask mode. Peak RSS grew only from 54.2 to 96.2 MiB in f64 for 5.4 times as many structures; it increased with the number of workers (Table S7) and not with the size of the collection. The comparators used 3.4–7.2 times as much (Table S8).

In a larger single run, we processed the 550,122 Swiss-Prot structures of the AlphaFold Database (PDB format, f32, 128 sphere points; one measured run per condition). With 10 workers, zsasa processed the collection at 1,599 structures/s in bitmask f32 mode (5.7 min) and at 735 structures/s in exact f32 mode, with 291 and 289 MiB peak RSS (Figure S3; Table S7). Lahuta’s bitmask mode, measured at 10 threads only, processed it at 1,269 structures/s with 660 MiB peak RSS. The 10-worker bitmask run kept 4.8 of the 10 cores busy on average, because each file that is opened for the first time waits for the on-access file scanner of the test machine. Supplementary Note S2 describes this effect and the disk I/O policy used for these runs, and reports runs with more workers than cores.

#### Large and parser-heavy single structures

Single-file stress tests reveal parser effects and large-structure behavior that directory-level summaries can hide. The single-file subset contains eight structures from the AlphaFold Database and the PDB, ranging from 10,919 to 4,506,416 atoms and from 1 to 1,574 chains, read as mmCIF (Table S9). Runtime increased smoothly with atom count, and the f64 mode was 2.4-to 5.4-fold faster than FreeSASA and 1.4-to 3.8-fold faster than RustSASA across the size range, with peak RSS lower by factors of 8.3–17.0 and 3.1–7.1 (Figure 4a,b; Lahuta cannot process most of these inputs). The largest case, a hetero 1,574-mer with 4,506,416 atoms, completed in 6.266 s (f64) and 5.324 s (bitmask f32) at 100 points and 10 threads, whereas FreeSASA, RustSASA, and PDBTools.jl (Martínez and Lima 2026) required 28.95 s, 23.65 s, and 10.51 s, corresponding to 4.62-fold, 3.77-fold, and 1.68-fold speedups for the f64 mode. Component timing attributed 2,015 ms to parsing and 3,770 ms to SASA calculation in the f64 run, and the bitmask run reduced the SASA component to 2,846 ms (Figure 4c); parsing took 20.0 s in FreeSASA and 16.4 s in RustSASA. The bitmask mode was slower than the exact mode below about 100,000 atoms because of the fixed cost of building its lookup table.

**Figure 4.**
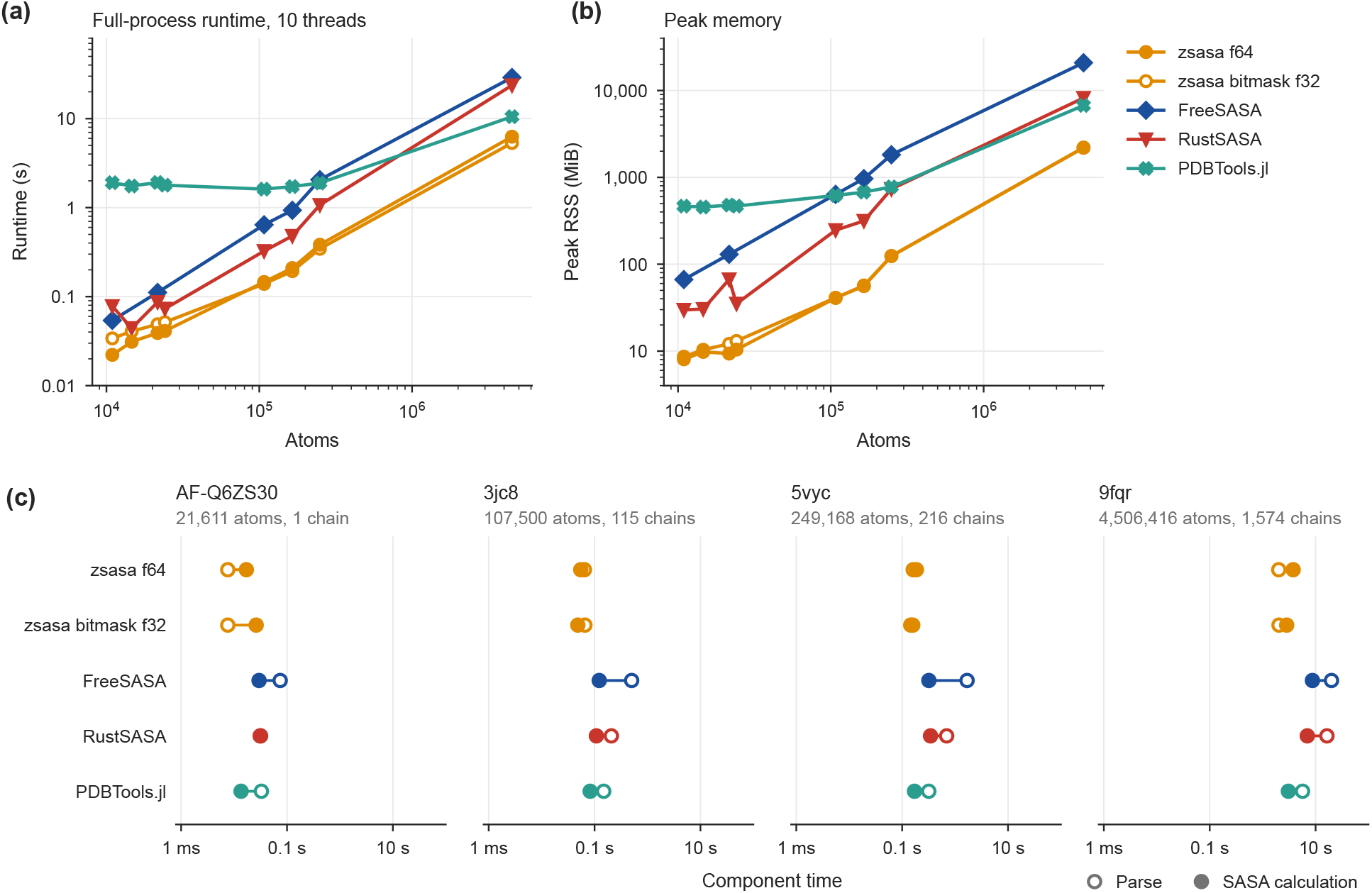
Single-structure runtime, peak memory, and component timing (mmCIF input, 10 threads, 100 sphere points). (a) Full-process runtime and (b) peak RSS versus atom count for the eight structures of the single-file subset (Table S9). FreeSASA has no values for the two AFDB-derived complexes, and Lahuta is excluded because it cannot process most of these inputs. (c) Parse time (open markers) and SASA-calculation time (filled markers) for four structures of increasing size. PDBTools.jl full-process runtime includes Julia startup, whereas its component timings come from a warm process. The remaining component timings are provided in Figure S4.

We used mmCIF for this suite because the PDB format cannot represent the largest assemblies faithfully. Coordinates above 999.999 Å fill the fixed-width coordinate fields completely, and FreeSASA then misreads adjacent fields without reporting an error. On PDB versions of 8rbs and of 9fqr, the largest case, FreeSASA took 14.0 s and 199.5 s, against 0.93 s and 28.95 s on mmCIF. RustSASA likewise spent 13.0 s parsing the PDB version of 5vyc, against 1.07 s in total on mmCIF. Per-structure results are provided in Table S10.

### Trajectory benchmarks

#### Validation against MDTraj

For trajectories, we compared per-frame total SASA with that from MDTraj on the 5wvo_C trajectory (1,001 frames; Figure 5a). When `zsasa` was called through its MDTraj integration, which uses MDTraj’s element radii, it agreed with MDTraj without bias: the median signed relative difference was within 0.02% at every sphere-point count, and the frame-to-frame scatter fell from a mean relative difference of 0.38% at 64 points to 0.05% at 1,024 points (residual *R*^2^ from 0.973 to 0.9995). The remaining scatter is consistent with sphere-point placement along a different axis in MDTraj (Table S11). The CLI f64 mode differed from MDTraj by a nearly constant −0.17% to −0.20% across point counts, which reflects the different radii: NACCESS radii for the CLI and MDTraj’s own element radii. The CLI bitmask mode differed by a constant −2.0%, a larger offset than for static structures. We therefore do not recommend the bitmask mode for explicit-hydrogen trajectories; an experimental bias-correction option that removes most of this offset is documented in Table S12. During this validation we found that MDTraj returns inflated areas for every frame after the first when a multi-frame trajectory is passed to its Shrake–Rupley function in a single call, by 1.5% at 64 points and 0.09% at 1,024 points for this trajectory. The reference values were therefore computed one frame at a time. Full results are given in Tables S12 and S13.

**Figure 5.**
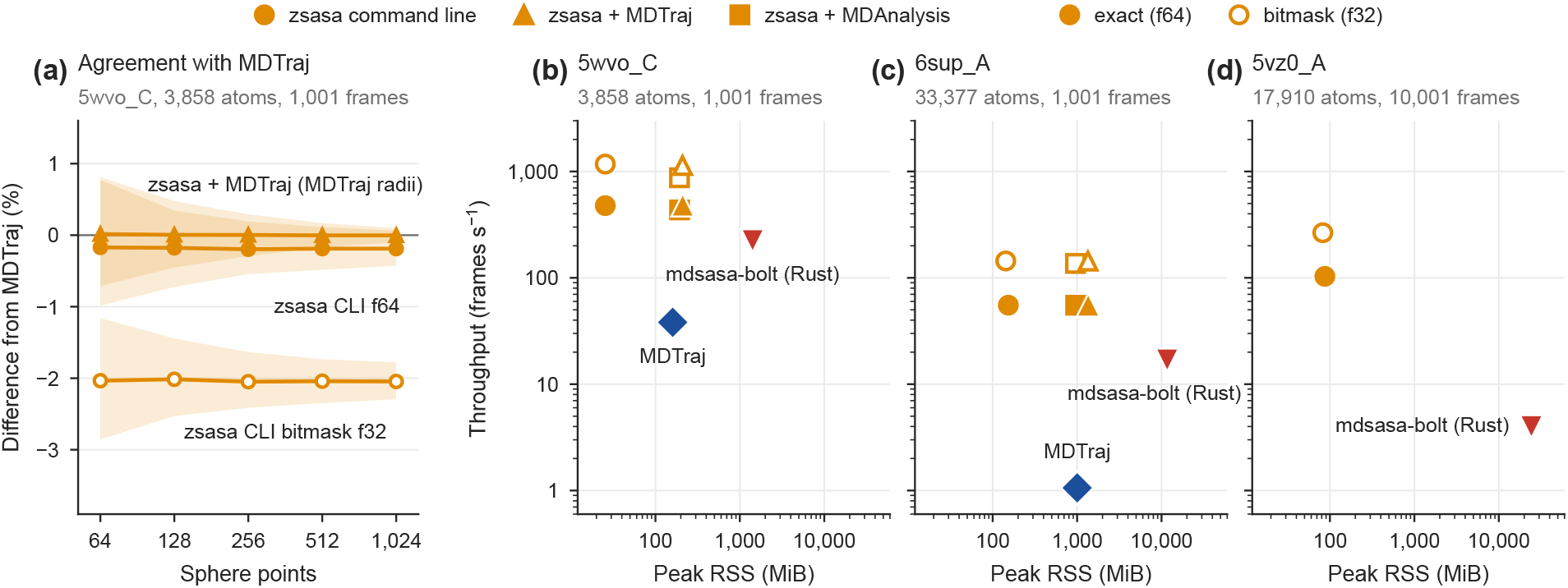
Trajectory agreement with MDTraj, throughput, and peak memory. (a) Signed relative difference from MDTraj on the 5wvo_C trajectory (1,001 frames) for zsasa called through its MDTraj integration (MDTraj radii) and for the CLI in f64 and bitmask f32 modes (NACCESS radii). Lines show the median and shaded bands span the 5th–95th percentiles; negative values indicate lower total SASA than MDTraj. (b–d) Throughput versus peak RSS at 128 sphere points and 10 threads for (b) 5wvo_C, (c) 6sup_A, and (d) 5vz0_A on shared logarithmic axes; higher and farther left is better. Marker shape distinguishes the `zsasa` CLI from its MDTraj and MDAnalysis integrations, and filled and open markers denote the exact f64 and bitmask f32 modes. Throughput values are means of three runs. Neither MDTraj nor the Python integrations were run on the 10,001-frame 5vz0_A trajectory.

#### Trajectory throughput

At 128 sphere points and 10 threads, the CLI f64 mode processed the 5wvo_C, 6sup_A, and 5vz0_A workloads in 2.098 s, 18.098 s, and 96.858 s, corresponding to 477, 55.3, and 103 frames/s, with peak RSS between 25.5 and 153.2 MiB. zsasa again combined the highest throughput with the lowest peak RSS (Figure 5b–d), and its Python integrations reached similar throughput (100–101% of the CLI value with MDTraj and 91–100% with MDAnalysis) plus the memory used by the host library’s trajectory reader. MDTraj’s own Shrake–Rupley function ran at about 38 frames/s on 5wvo_C and about 1 frame/s on the larger 6sup_A system. mdsasa-bolt (Rust, Campbell 2026) used far more memory because its MDAnalysis front end loads the atom data for every frame before the Rust core runs, not because of the SASA calculation. The f64 mode was 12.5-fold and 52.2-fold faster than MDTraj on 5wvo_C and 6sup_A, and 25.9-fold faster than mdsasa-bolt on 5vz0_A (Table 3; Table S14). The bitmask mode was a further 2.5-to 2.6-fold faster (1,174, 144, and 266 frames/s) with the same memory use, but with the −2.0% offset described above.

### Large-scale workflow demonstration

As an end-to-end demonstration, we computed interface burial for two sets of predicted dimers released as part of the AlphaFold Database complex-structure expansion: the 91,637 human homodimer models and 531,605 heterodimer models in which both chains are human proteins (Miller et al. 1987; Janin, Miller, and Chothia 1988; Schweke et al. 2024; Han et al. 2026). We treat these datasets as a workflow case and not as a performance benchmark, because they combine requirements that the comparator tools do not uniformly cover: zstd-compressed mmCIF inputs, chain-aware analysis of two partners, and integration with external confidence annotations. For each model, a two-partner workflow reported the SASA of chain A, chain B, and the complex, and their difference, ΔSASA = SASA_*A*_ + SASA_*B*_ − SASA_*AB*_, which is twice the buried surface area per chain (Chakravarty et al. 2013) and is the form commonly used as an interface descriptor in binder design (Overath et al. 2025).

One batch command per set, in bitmask f32 mode at 128 sphere points with 20 workers on the 10-core laptop used for the benchmarks, processed the homodimers in 2 min 57 s and the heterodimers in 23 min 39 s. All 623,242 models were processed without failure, with peak RSS of 1,655 and 350 MiB, respectively (Table S15). Each set was run once with `zsasa` v0.12.1 under the default disk I/O policy of macOS; a second run of each under the passive disk I/O policy (Supplementary Note S2) changed the wall time by about 1% or less and gave the same results.

The two sets had similar interface sizes (Figure 6a): the median ΔSASA was 2,829 Å^2^ (interquartile range 1,807–4,536 Å^2^) for homodimers and 2,933 Å^2^ (1,793–4,641 Å^2^) for heterodimers. ΔSASA was zero for 1.0% and 1.7% of the models and never negative.

**Figure 6.**
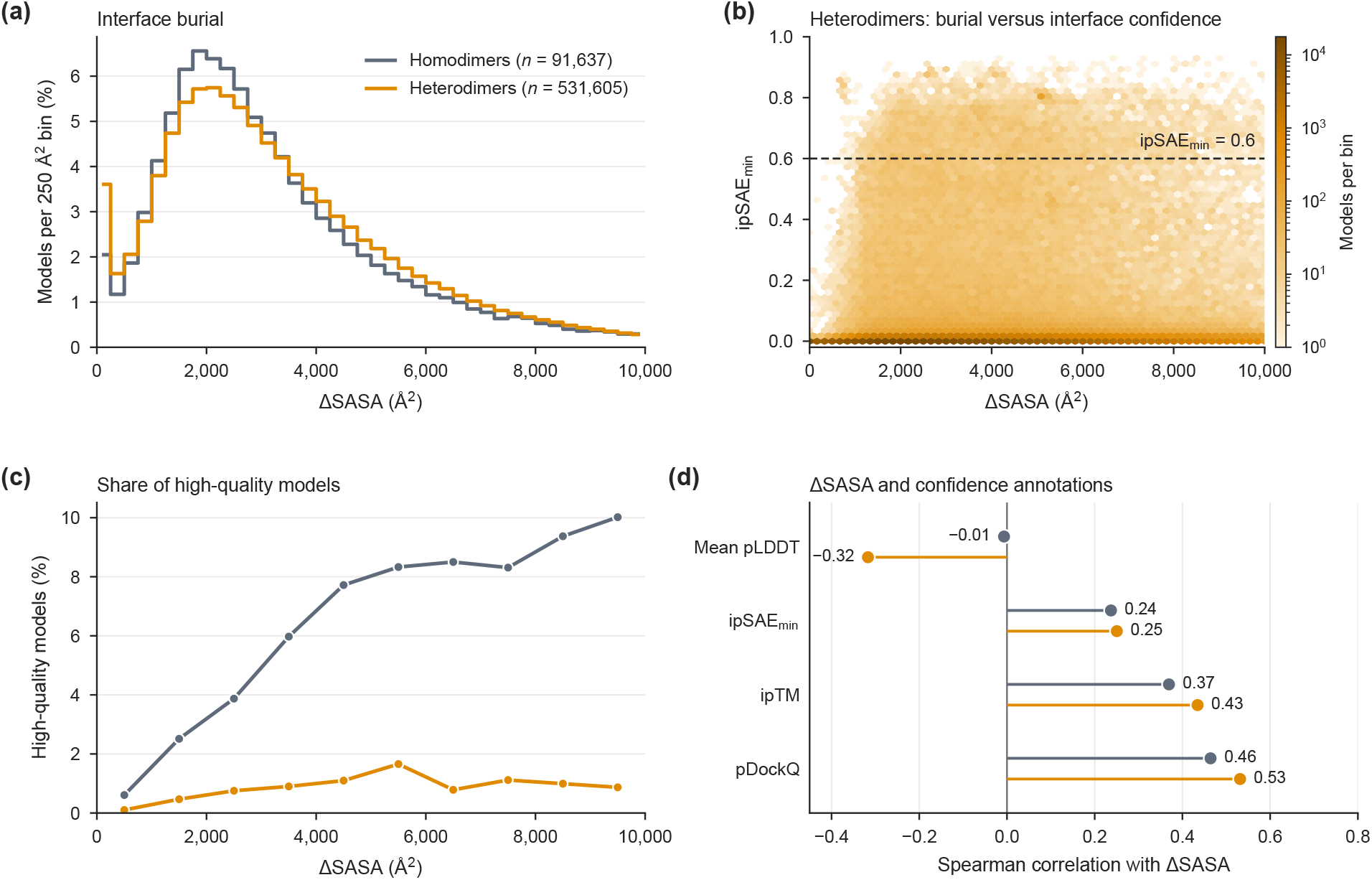
Interface burial in AFDB human dimer models. ΔSASA from `zsasa` for 91,637 human homodimer models and 531,605 human–human heterodimer models. (a) Distribution of ΔSASA in 250 Å^2^ bins; 4.9% of homodimers and 2.6% of heterodimers lie beyond 10,000 Å^2^. (b) ΔSASA versus ipSAE_min_ for the heterodimers, shown as model density; the dashed line marks the ipSAE_min_ = 0.6 threshold. (c) Share of high-quality models in 1,000 Å^2^ bins of ΔSASA. (d) Spearman correlation between ΔSASA and four confidence annotations. Gray, homodimers; orange, heterodimers.

Because these are predicted complexes, burial should not be read as standalone evidence for an interaction, so we compared it with the AFDB confidence annotations. Following the AFDB definition (Supplementary Note S3), models were labeled high quality when ipSAE_min_ ≥ 0.6, mean pLDDT ≥ 70, and backbone clashes ≤ 10 (Han et al. 2026); 5.1% of the homodimers and 0.8% of the heterodimers met these criteria, and ipSAE_min_ was zero for 53.1% and 79.6% of the models. A large buried surface did not imply a confident interface (Figure 6b). Among homodimers, the share of high-quality models rose from 0.6% below 1,000 Å^2^ to about 8% at 4,000–5,000 Å^2^ and more slowly beyond that (10% above 9,000 Å^2^), whereas among heterodimers it stayed below 2% at every interface size (Figure 6c). Rank correlations between ΔSASA and the confidence annotations were moderate for pDockQ (Spearman ρ = 0.46 and 0.53 for homo- and heterodimers) and ipTM (0.37 and 0.43), weak for ipSAE_min_ (0.24 and 0.25), and absent or negative for mean pLDDT (−0.01 and Confidence scores summarize model-level support for a predicted structure, whereas ΔSASA quantifies the geometric extent of an interface; the two are complementary, as a recent meta-analysis of de novo binder designs also found (Overath et al. 2025). This analysis is a scale demonstration and not a biological discovery claim. The per-model values are released as tables that can be searched and filtered in the browser on the documentation site (Data availability).

## Discussion

The exact mode of `zsasa` closely matches FreeSASA under matched settings, so its values are compatible with existing Shrake–Rupley-based analyses, and the systematic offset of the approximate bitmask mode has been measured. `zsasa` is nevertheless more than a Zig rewrite of an existing SASA calculator. The computational core uses standard algorithms, but the tool combines compressed input, directory-scale batch processing, streaming JSON Lines output, residue-level mappings, Python access, and trajectory input in one program, and the same engine handled inputs from one structure to several hundred thousand. For large datasets, users also need outputs that can be consumed incrementally, failures that can be isolated to individual structures, and tables that can feed downstream statistical or machine-learning analyses. Point-by-point comparison with existing tools also proved useful beyond validation by exposing the inflated multi-frame areas of MDTraj.

The current implementation and evaluation have several limitations. First, Zig remains pre-1.0, so the implementation may require maintenance as the language, standard library, and build system evolve. Second, `zsasa` is still an early-stage project, and its command-line options, Python API, and output formats may change. Third, the CLI intentionally performs only limited structure preparation; alternate-location policy, complex cleanup, specialized selections, and input normalization should be handled in Python or in upstream preprocessing. Fourth, the bitmask mode is approximate and is not recommended for explicit-hydrogen trajectories, so analyses that depend on small absolute differences or residue-level values should use the exact mode or include independent validation.

The benchmarks should also be read within their scope. All head-to-head timings were collected on one laptop and operating system, with pinned comparator builds and specific point counts; different CPUs, storage systems, compiler settings, or workload mixtures may change absolute runtimes and some speedup ratios. Trajectory validation was performed on one ATLAS trajectory. At the largest scale tested, throughput also depended on how quickly input files could be opened: the Swiss-Prot run kept peak RSS below 600 MiB for more than half a million structures (Figure S3), while about half of the cores were idle, waiting for file access (Table S7; Supplementary Note S2).

Standard SASA calculations can now be run over entire predicted-structure collections on a single laptop. zsasa remains under active development, and we will continue to adapt its performance, input handling, outputs, and workflow options to the analyses it is used for.

## Materials and Methods

### Implementation

#### Algorithms

The core Zig library separates parsing, atom classification, SASA algorithms, compression, batch execution, trajectory handling, and output serialization. The Shrake–Rupley (Shrake and Rupley 1973) and Lee–Richards (Lee and Richards 1971) paths follow the original algorithms (Results). The bitmask mode bins directions on an octahedral grid, batches neighbor tests with explicit SIMD vectors, and combines occlusion masks with bitwise AND and population-count operations.

#### Atom classification

Radii can be supplied directly or assigned through the CCD, ProtOr, NACCESS, and OONS-style classifiers (John D. Westbrook et al. 2015; Hubbard and Thornton 1993; Ooi et al. 1987; Tsai et al. 1999). For standard residues, the default CCD classifier uses a compiled ProtOr-compatible table of united-atom radii (Tsai et al. 1999). For other components, it derives ProtOr-compatible radii and polarity classes from the atom and bond tables of the wwPDB Chemical Component Dictionary, following the atom-typing logic described in the ChimeraX documentation, in which united-atom radii depend on atom type, hybridization, and the number of attached hydrogens (UCSF Resource for Biocomputing, Visualization, and Informatics 2020; John D. Westbrook et al. 2015). Unsupported atoms fall back to element-based radii (Bondi 1964; Mantina et al. 2009). Custom radius tables can be supplied as TOML classifier files.

#### Interfaces and formats

The CLI detects the input format (PDB, mmCIF, BinaryCIF, or SDF/MOL) and reads gzip- and zstd-compressed files directly (wwPDB Consortium 2019; John D. Westbrook et al. 2022; Sehnal et al. 2020). zsasa can also be called from Python, where it adds relative solvent accessibility and residue-level analysis (Tien et al. 2013) and integrates with Gemmi, Biopython, Biotite, MDTraj, and MDAnalysis (Wojdyr 2022; Cock et al. 2009; Kunzmann and Hamacher 2018; Kunzmann et al. 2023; McGibbon et al. 2015; Michaud-Agrawal et al. 2011; Gowers et al. 2016).

#### Performance engineering

Coordinate, radius, and intermediate arrays are stored in compact contiguous buffers, temporary storage is scoped to the structure, batch job, or trajectory frame being processed, and directory runs stream inputs and outputs rather than retaining whole collections in memory. The kernels use Zig’s explicit @Vector operations, which Zig/LLVM lowers to the target instruction set, with a compile-time check enabling a wider AVX-512F path when available. Parallelism depends on the mode. For a single structure, atom ranges are distributed among threads through a shared atomic counter. In batch mode, each worker thread processes whole files, and the number of workers may exceed the number of CPU cores. In trajectory mode, frames are processed in bounded batches and results are written in frame order. Dependencies and the build are described in Supplementary Note S1.

#### Workflows and outputs

Results are written as JSON, JSON Lines, or CSV; the workflow and output options are described in the user documentation (https://n283t.github.io/zsasa/).

### Benchmark design

All head-to-head comparisons used zsasa v0.12.1 and the pinned comparator builds and Python dependencies listed in Table S16; comparator builds were pinned with Nix, and the Python environment was managed with uv (Python 3.12). Validation, batch, and trajectory runs were repeated for all tools in the same session as zsasa; the single-file comparator timings and the mdsasa-bolt timing on 5vz0_A were retained from earlier sessions on the same machine. Benchmarks were run on a MacBook Pro with an Apple M4 processor (10 cores: 4 performance and 6 efficiency) and 32 GB of memory, running macOS 26. Wall-clock runtimes are means of three runs after warmup runs (Table S17), which leave the input files in the file cache, so the three-run timings are warm-cache results. The Swiss-Prot run was timed once without warmup: its files are too many to stay in the file cache, or among the files recently checked by the on-access scanner of the test machine, between runs. Its commands were run under the passive disk I/O policy of macOS (`taskpolicy -d passive`), for `zsasa` and Lahuta alike, so that a tool’s own reads did not throttle the scanner (Supplementary Note S2). Peak memory is reported as peak RSS, and speedup ratios are comparator runtime divided by `zsasa` runtime. Timing tools, run-to-run variation, and the separation of parse and SASA-calculation times are described in Supplementary Note S4.

Unless stated otherwise, throughput benchmarks used 10 threads. Batch and trajectory throughput runs used 128 sphere points, a count supported by Lahuta’s bitmask mode, and single-file runs used 100, the FreeSASA default. Validation runs covered sphere-point counts from 64 to 1,024. Trajectory benchmarks used explicit hydrogens at stride 1 on all-atom trajectories from the ATLAS database (Vander Meersche et al. 2024), and the CLI runs used NACCESS radii. Except for these CLI runs, all tools used their default radii and probe radius (1.4 Å). The MDTraj (1.11.1) reference was computed one frame at a time (Results; reported at https://github.com/mdtraj/mdtraj/issues/2195); MDTraj throughput was timed with the usual single call. The exact commands are in the benchmark repository. Validation is a consistency check against existing tools under matched settings, not a comparison with an external ground truth, because SASA is defined operationally by the algorithm, radii, probe radius, sampling convention, hydrogen policy, and parser decisions.

Several comparator details affect how the results should be read. FreeSASA has no native directory mode, so the FreeSASA batch numbers were produced with `freesasa_batch`, a thin multi-file wrapper that we built around the pinned FreeSASA library. Lahuta’s SASA command requires AlphaFold2 models as input and processes only chain A, so it appears only in the validation and batch comparisons. PDBTools.jl, a Julia package with a Shrake–Rupley implementation (Martínez and Lima 2026), was included in the single-file comparison. Further details, including output granularity, single-file input preparation, and the Lee–Richards check, are given in Supplementary Note S4 and Tables S1, S9, S11, and S16–S19.

### AFDB dimer workflows

Structure files and score metadata were taken from the AFDB/NVIDIA FTP release (https://ftp.ebi.ac.uk/pub/databases/alphafold/collaborations/nvda/). The homodimer set comprises the human models with available structure files, and the heterodimer set the 531,605 models in which both chains are human proteins. Both runs used the default CCD classifier and a 1.4 Å probe with `zsasa` v0.12.1. Input preparation and derived quantities are described in Supplementary Note S3.

## Supplementary Material

The Supplementary Material contains Supplementary Notes S1–S4, Tables S1–S19, and Figures S1–S4: benchmark settings and environment, full validation and performance tables, implementation and methodology details, and a summary of the AFDB dimer workflows.

## Acknowledgments

This work relies on open-source structural-biology software, including FreeSASA, RustSASA, Lahuta, MDTraj, and MDAnalysis, both as comparators and as components (Mitternacht 2016; Campbell 2026; Sejdiu and Babu 2026; McGibbon et al. 2015; Michaud-Agrawal et al. 2011). We thank the developers and maintainers of these projects. This work was supported by JST, NBDC Grant Number JPMJND2401, Japan (T.N. and K.T.), and was partially supported by the Research Support Project for Life Science and Drug Discovery (Basis for Supporting Innovative Drug Discovery and Life Science Research (BINDS)) from AMED under Grant Number JP26ama121028 (K.T.).

## Author contributions

**Tsubasa Nagae:** Conceptualization; Methodology; Software; Validation; Formal analysis; Investigation; Data curation; Visualization; Writing – original draft. **Kentaro Tomii:** Supervision; Funding acquisition; Writing– review & editing.

## Conflict of interest

The authors declare no conflicts of interest.

## Data availability

zsasa is open-source software released under the MIT license. Source code is available at https://github.com/N283T/zsasa, and user documentation, including installation instructions and examples, is hosted at https://n283t.github.io/zsasa/. The Python package is distributed through PyPI (https://pypi.org/project/zsasa/) with pre-built wheels for Linux, macOS, and Windows and can be installed with `pip install zsasa`. The software version evaluated in this manuscript is `zsasa v0.12.1`.

Benchmark code, run configurations, result tables, and figure-reproduction materials are maintained in the `zsasa-benchmarks` repository. The repository also holds the per-run timings and their standard deviations, the full thread-count sweeps, further summary figures, and the investigation of file-access latency in the Swiss-Prot run (`investigations/swissprot-open-latency`). The release corresponding to `zsasa` v0.12.1 is available at https://github.com/N283T/zsasa-benchmarks/releases/tag/v0.12.1-benchmark-archive-r3 and is deposited in Zenodo at https://doi.org/10.5281/zenodo.23898582.

The AlphaFold Database structures used in the static-structure benchmarks were obtained from the AlphaFold Database portal (https://alphafold.ebi.ac.uk/) and FTP area (https://ftp.ebi.ac.uk/pub/databases/alphafold/), with the exact dataset manifests and filters recorded in the benchmark repository. The molecular-dynamics trajectory benchmarks used all-atom trajectories obtained from the ATLAS database (https://www.dsimb.inserm.fr/ATLAS/index.html) (Vander Meersche et al. 2024), with trajectory identifiers and run settings recorded in the benchmark repository and the Supplement.

The AFDB/NVIDIA dimer source structures and score metadata used in the workflow demonstration were obtained from the AFDB/NVIDIA FTP release at https://ftp.ebi.ac.uk/pub/databases/alphafold/collaborations/nvda/. These large upstream files are not redistributed here. The workflow files, run records, and per-model ΔSASA values of the two dimer runs are included in the benchmark repository (`workflows/afdb-dimers/`). Per-model tables of the dimers and of the 550,122 AFDB Swiss-Prot models, computed with zsasa v0.13.0 (which adds the polar/apolar split and gives the same totals), are in the same archive and can be searched and downloaded through the SASA explorer of the documentation site (https://n283t.github.io/zsasa/docs/data/explorer/).

## AI disclosure

AI-assisted tools, including OpenAI ChatGPT/Codex and Anthropic Claude/Claude Code, were used to help write and debug code, organize the benchmarks, draft the manuscript, and edit its language. The authors conceived and designed the study, chose the analyses, interpreted the results, made all final decisions on the manuscript, and reviewed and edited all AI-assisted output.

## Notes

https://github.com/N283T/zsasa

https://doi.org/10.5281/zenodo.23898582

https://github.com/N283T/zsasa-benchmarks/releases/tag/v0.12.1-benchmark-archive-r3

https://n283t.github.io/zsasa/

https://pypi.org/project/zsasa/

### Competing Interest Statement

The authors have declared no competing interest.

### Summary of Updates

Benchmarks rerun with zsasa v0.12.1 (previously v0.6.0); all values, figures, and tables updated. Manuscript restructured and condensed; figures redrawn; Supplementary Material reorganized. New results: per-atom and per-residue agreement with FreeSASA, a Swiss-Prot run of 550,122 structures, PDBTools.jl as an additional comparator, and an MDTraj multi-frame issue reported upstream. Workflow demonstration extended to 531,605 human heterodimer models in addition to 91,637 homodimers. Benchmark data deposited in Zenodo, with per-model tables searchable online.

