## Supplementary Material for "zsasa: a Zig-based engine for high-throughput solvent accessible surface area at proteome scale"

The Supplementary Notes, Tables, and Figures are grouped in that order, and each group is numbered in the order of first citation in the main text. The table below lists, for each section of the main text, the supplementary items that support it. Per-run records, the full thread-count sweeps, further summary figures, and the logs of the investigations are in the benchmark archive (Data availability in the main text).

| Main-text section | Supplementary material |
|---|---|
| Design of `zsasa`; Materials and Methods, Implementation | Supplementary Note S1 |
| Benchmark overview; Materials and Methods, Benchmark design | Tables S1, S11, and S16–S19; Supplementary Note S4 |
| Validation against FreeSASA | Tables S2–S4; Figures S1a and S2 |
| Proteome-scale batch throughput | Tables S5–S8; Figure S3; Supplementary Note S2 |
| Large and parser-heavy single structures | Tables S9 and S10; Figure S4 |
| Validation against MDTraj | Tables S11–S13; Figure S1b |
| Trajectory throughput | Table S14 |
| Large-scale workflow demonstration; Materials and Methods, AFDB dimer workflows | Table S15; Supplementary Note S3 |

#### Supplementary notes

##### Supplementary Note S1. Implementation details and the role of Zig.

This note adds detail to the implementation summary of the main text (Design of zsasa; Materials and Methods). Atom classification, input formats, the Python API, and workflow files are described in the user documentation (https://n283t.github.io/zsasa/).

**Algorithms and approximate mode.** The exact mode computes Shrake–Rupley SASA (Shrake and Rupley 1973), and Lee–Richards (Lee and Richards 1971) provides an independent slice-integration path. The lookup-table design of the bitmask mode was inspired by Lahuta (Sejdiu and Babu 2026), but `zsasa` implements it independently rather than as a port: it accepts any point count from 1 to 1,024 rather than a fixed set of values, batches neighbor tests with explicit SIMD vectors, and bins directions on an octahedral grid. Because the bitmask path is an acceleration strategy rather than an identical numerical formulation, its agreement with reference outputs is reported separately from that of the exact mode.

**Performance engineering.** The compact buffers, the scoping of temporary storage to one structure, batch job, or frame, and the streaming of directory inputs and outputs (Materials and Methods) account for the low peak RSS in the batch and trajectory benchmarks, where the limiting resources are often input/output and transient working memory rather than floating-point throughput. Besides the optional AVX-512F path, the generic 8- and 4-wide vector paths are lowered by Zig/LLVM to the target instruction set, such as x86 AVX/SSE or ARM NEON, rather than through opportunistic autovectorization or hand-written per-ISA intrinsics. In batch mode, file-level workers claim work items atomically. Together with the atomic-counter distribution of atom ranges and the bounded frame batches, this keeps scheduling overhead small while allowing irregular workloads—structures with different atom counts, chain counts, or frame costs—to share one implementation style without requiring the entire dataset to be resident in memory.

**The role of Zig.** Although these optimizations are not unique to Zig and could be implemented in C or C++, Zig makes allocator choice, explicit SIMD, compile-time feature selection, and cross-platform build behavior visible in the source and build system. The SASA core uses only the Zig standard library, while trajectory-format readers come from `ztraj`, a first-party Zig module spun out of this project. This keeps the dependency footprint small and controlled. Zig's build system produces statically linked binaries and cross-compiles to Linux, macOS, and Windows from one toolchain, keeping installation and reproducible benchmarking straightforward. The comparators range from a C/C++ Autotools build with bundled and optional libraries (FreeSASA), through Rust crate dependencies (RustSASA), to more than a dozen bundled C++ libraries (Lahuta). Table 1 of the main text summarizes the SASA capabilities of each tool. Its dependency counts refer to direct dependencies in the compared SASA/CLI paths: `zsasa` counts `ztraj`; FreeSASA counts Gemmi, json-c, and libxml2; RustSASA counts direct Cargo dependencies with the CLI feature.

**Supplementary Note S2. File access in the Swiss-Prot run and worker counts above the CPU count.**

**On-access file scanning.** The test machine runs endpoint-security software with on-access scanning (Microsoft Defender for Endpoint), which checks a file when it is first opened and remembers recently checked files. Measured with one thread on 4,000 Swiss-Prot files, the first open of a file took 5.8 ms and a repeated open 0.2 ms; reading the file took 0.2–0.3 ms either time. In sampled runs on files that had not been scanned, the `zsasa` workers spent 64–71% of their wall time inside the open call. The three-run timings of the other suites were not affected, because their warmup run absorbs the scan: an 8,000-file subset that had been scanned ran equally fast under either disk I/O policy described below. The Swiss-Prot collection is too large for that. It exceeds the memory of the machine and, judging from repeated runs, what the scanner remembers, so every run of it opens files that are scanned again.

**Disk I/O policy.** macOS holds back lower-priority disk I/O while a normal-priority process is doing I/O on the same device, and the scanner appeared to be subject to this throttling: a batch tool slowed down the scan that its own workers were waiting for. Under the passive disk I/O policy (`IOPOL_PASSIVE`, set here by starting each command with `taskpolicy -d passive`), a process keeps its priority but no longer throttles lower-priority I/O. The Swiss-Prot results in the main text were obtained under this policy, for `zsasa` and Lahuta alike, and nothing else differed from the default-policy commands. The default-policy rows of Table S7 were measured with `zsasa` v0.12.1 and Lahuta in a separate session in October 2026. Under that policy both tools were slower (Table S7; Figure S3). At 10 workers, `zsasa` bitmask f32 was 12% faster than Lahuta's bitmask mode under the default policy and 26% faster under the passive policy.

**Earlier versions.** With `zsasa` v0.9.0, which memory-mapped its inputs and wrote more output, the order was reversed under the default policy: Lahuta was 14% faster at 10 workers. Since v0.11.0, `zsasa` reads each batch input with a single read and has a `--passive-io` option that requests the passive policy; the v0.12.1 runs of Table S7 nonetheless set the policy through `taskpolicy`, as for Lahuta. The v0.9.0 measurements, the analysis of memory mapping and the file cache, and the scripts and logs of the measurements in this note are archived with the benchmark repository (`investigations/swissprot-open-latency`).

**Worker counts above the CPU count.** In batch mode each worker thread processes one file at a time, and `zsasa` allows more workers than CPU cores. On the *E. coli* and Human collections, using more workers than cores changed throughput by less than 3% and increased peak RSS; on Swiss-Prot under the passive policy, 20 workers raised bitmask f32 throughput by 4% (1,666 structures/s) and 40 workers lowered it by 2% (1,566 structures/s), while peak RSS rose to 373 and 531 MiB, respectively (Table S7). The *E. coli* and Human collections occupy 0.8 and 9.1 GiB on disk, respectively, and fit in the file cache of the 32 GB test machine, and `zsasa` kept 8.6–9.6 of the 10 cores busy on them. On the 119 GiB Swiss-Prot collection, the bitmask f32 runs kept 4.8, 5.2, and 4.9 cores busy with 10, 20, and 40 workers, respectively.

**Supplementary Note S3. AFDB dimer workflow: derived quantities and inputs.** For a dimer model with chains A and B, $\Delta\text{SASA} = \text{SASA}_A + \text{SASA}_B - \text{SASA}_{AB}$, where $\text{SASA}_A$ and $\text{SASA}_B$ are computed for each chain in isolation and $\text{SASA}_{AB}$ for the complex. The buried surface area per chain is $\Delta\text{SASA}/2$. $\text{ipSAE}_{\text{min}}$ is the smaller of the two directional ipSAE values of a model. Following the AFDB complex-confidence definition, a model is labeled high quality when $\text{ipSAE}_{\text{min}} \geq 0.6$, mean pLDDT ≥ 70, and the number of backbone clashes is at most 10. Mean pLDDT is the average over all residues of a model. The homodimer set comprises 91,635 models in mmCIF and two in PDB format; four entries had no structure file in the source archives. The heterodimer models had been stored locally in a lossless compact encoding and were restored to zstd-compressed mmCIF before the run.

**Supplementary Note S4. Benchmark methodology details.**

**Scope.** The large-scale validation and performance suites focus on the Shrake–Rupley exact mode and the Shrake–Rupley bitmask approximation. Lee–Richards is treated as a secondary compatibility path: it was checked only at 20 slices against FreeSASA (Table S4), and that check should be read as limited supplemental validation rather than as a benchmark at the scale of the Shrake–Rupley analyses. Validation (suites 1 and 4) is a consistency check under matched settings rather than a comparison with an external ground truth (Materials and Methods). Close agreement shows that `zsasa` stays in line with established tools, not that any tool is uniquely “correct”. The implementation differences behind the remaining disagreements are listed in Table S11 and discussed with the validation results.

**Recorded fields.** The benchmark harness recorded the same fields for every run: input identity, tool and mode, point count or slice count, thread count, wall-clock timing where applicable, peak resident set size (RSS), and the output values needed for agreement or speedup calculations. Tables S17 and S18 summarize the five suites; the run configurations, manifests, and per-run records are in the benchmark archive (Data availability in the main text).

**Comparators.** Several comparator details affect how the results should be read. First, FreeSASA has no native directory or batch mode. Because invoking it sequentially over a proteome-scale directory is prohibitively slow, the FreeSASA batch numbers were produced with `freesasa_batch`, a thin multi-file wrapper we built around the C API of the pinned FreeSASA library specifically for these benchmarks, so that all tools are timed as multi-file workloads rather than as shell-level loops. In the batch results, “FreeSASA” therefore refers to this wrapper, not to a native FreeSASA feature. Second, Lahuta appears only in the validation and batch comparisons: its SASA command requires AlphaFold2-model input and hardcodes chain “A”, so it cannot process the multi-chain and large-assembly structures in the single-file subset and is excluded there. Third, the single-file comparison additionally includes PDBTools.jl (Martínez and Lima 2026), a Julia package whose full-process runtime includes Julia startup; its component timings were measured in a warm process and exclude that startup. Fourth, the tools differ in batch output granularity: `zsasa` and Lahuta write per-atom areas, RustSASA per-residue values, and FreeSASA only the total. `zsasa` emits the most detailed output of the four, so its batch-throughput advantage is, if anything, conservative; we did not separately quantify this output cost.

**Timing.** Wall-clock runtimes were recorded with hyperfine 1.20.0, with a filesystem sync between runs, and per-run records were aggregated in DuckDB. The pinned comparators did not change between `zsasa` releases. The standard deviation across the three runs was at most 1.0% of the mean for the 10-thread batch timings, except for RustSASA (16.6% on the *E. coli* collection and 2.5% on the Human collection), and at most 3.3% for the trajectory timings; per-run standard deviations are included in the result tables of the benchmark archive. Parse and SASA-calculation times were separated with a `--timing` option that is native to `zsasa` and that we added to the FreeSASA and RustSASA builds.

**Static validation.** Static validation tested whether `zsasa` reproduces FreeSASA total SASA under matched Shrake–Rupley settings. We used the *E. coli* AlphaFold Database collection because it provides thousands of independent predicted structures while remaining small enough for repeated point-count sweeps. Agreement was also checked at finer levels for the f64 mode at 128 points: the 10,520,167 per-atom areas and 1,349,634 per-residue areas of the 4,370 structures were compared with the atom- and residue-level text output of the FreeSASA command-line program, which carries two decimal places, and none differed at that precision (maximum absolute difference 0.005 Å$^2$). On the same structures and under matched settings, the 20-slice Lee–Richards run of `zsasa` reproduced the FreeSASA totals as closely as the Shrake–Rupley runs did (Table S4; Figure S2).

**Batch throughput.** Batch benchmarks measured complete directory processing, including parsing, SASA calculation, output writing, and worker scheduling. The *E. coli* collection was used for both comparator benchmarking and thread scaling, the Human collection tested the same 10-thread setting on a larger proteome, and the Swiss-Prot collection was processed once per condition (Supplementary Note S2).

**Single-file inputs.** Single-file benchmarks focused on large individual structures and on parsing cost. The subset (Table S9) spans a grid of sizes and chain counts up to the largest PDB-derived assembly, plus two comparator stress cases: 5vyc, a RustSASA parser outlier, and 8rbs, a FreeSASA runtime outlier. These two represent a broader set of structures that earlier benchmarking (in the `zsasa` repository) flagged as comparator outliers with an automatic per-bin detector; the others were not re-measured under the present pinned conditions, but the comparator parsers are unchanged, so the same behavior is expected to persist. All tools were timed on the same uncompressed mmCIF files so that differences reflected the benchmarked tool behavior rather than incompatible input handling; 5vyc and 9fqr were reduced to their protein atoms (221,406 and 20,949 other atoms removed), and the other six structures were used as deposited. FreeSASA stopped without assigning atomic radii on the two AFDB-derived complexes and has no values for them.

An earlier version of the suite used protein-only PDB inputs instead. Each source structure had hydrogens, alternative conformations, ligands, waters, and non-L-peptide chains removed, with residue numbers and atom serials wrapped into the PDB field limits and the `CRYST1` Z field filled. This normalization was required by the comparators rather than by `zsasa`: RustSASA's pdbtbx-based reader fails on the raw hybrid36 and oversized-field records that large assemblies produce, so without it several structures could not be parsed at all. `zsasa` reads only the records its SASA calculation needs and accepts the raw inputs directly; all tools were nonetheless run on the same normalized PDB so that the timing reflected SASA throughput rather than differing parser behavior. The suite moved to mmCIF because PDB versions of the largest assemblies cannot be represented faithfully: coordinates above 999.999 Å fill the fixed-width coordinate fields, which FreeSASA then misreads, and one-character chain identifiers must be reused. On the PDB inputs at 10 threads, FreeSASA took 14.0 s with 23.8 GiB peak RSS on 8rbs and 199.5 s with 23.8 GiB on 9fqr, and RustSASA took 13.3 s on 5vyc, of which 13.0 s was parsing.

**Trajectories.** Trajectory validation compared frame-wise total SASA against MDTraj as the reference; its goal was to measure convergence with point count, not to benchmark runtime. Trajectory-throughput benchmarks measured full trajectory processing, including trajectory reading and frame-wise SASA calculation. The largest trajectory used a reduced tool set to avoid impractically slow native MDTraj and Python-integration runs.

### Supplementary tables

**Table S1. Timing, aggregation, and derived metrics.**

| Quantity | Definition or policy |
|---|---|
| Runtime | Mean wall-clock seconds over the measured runs; warmup runs excluded |
| Runtime uncertainty | hyperfine standard deviation, kept in the benchmark archive |
| Peak RSS | Peak resident set size of each run, averaged over runs, MiB |
| Throughput | Item count (structures or frames) divided by mean runtime |
| ms/item | 1,000 × runtime divided by item count |
| Throughput/MiB | Throughput divided by peak RSS |
| Speedup | Comparator runtime divided by `zsasa` runtime |
| RSS reduction | Comparator RSS divided by `zsasa` RSS |
| Parallel speedup | Single-thread runtime divided by multithread runtime |
| Parallel efficiency | Parallel speedup divided by thread count |
| CPU proxy (busy cores) | User plus system time divided by wall time |
| Validation pairs | Shared records only; mean differences unweighted across structures or frames |
| Residual $R^2$ | $1 - SS_{\text{res}}/SS_{\text{tot}}$ versus the reference; absolute-agreement metric used for headline validation rows |
| Pearson $r^2$ | Squared Pearson correlation coefficient; co-variation metric reported for trajectory validation |
| Signed difference | Tool SASA − reference SASA |
| Relative difference | \|tool SASA − reference SASA\| / reference SASA × 100%; mean and maximum over structures or frames |
| ΔSASA | $\text{SASA}_A + \text{SASA}_B - \text{SASA}_{AB}$ |

**Table S2. Static validation, exact Shrake–Rupley modes.** Total SASA against FreeSASA on 4,370 *E. coli* AlphaFold Database structures (`zsasa` v0.12.1).

| Tool / mode | Points | Residual $R^2$ | Mean rel. diff. (%) | Max rel. diff. (%) |
|---|---:|---:|---:|---:|
| `zsasa` f64 | 64 | 1.000000 | 0.0000208 | 0.000221 |
| `zsasa` f64 | 128 | 1.000000 | 0.0000210 | 0.000227 |
| `zsasa` f64 | 256 | 1.000000 | 0.0000208 | 0.000257 |
| `zsasa` f64 | 512 | 1.000000 | 0.0000209 | 0.000245 |
| `zsasa` f64 | 1,024 | 1.000000 | 0.0000207 | 0.000205 |
| `zsasa` f32 | 64 | 1.000000 | 0.000108 | 0.0280 |
| `zsasa` f32 | 128 | 1.000000 | 0.000101 | 0.0153 |
| `zsasa` f32 | 256 | 1.000000 | 0.0000865 | 0.00790 |
| `zsasa` f32 | 512 | 1.000000 | 0.0000961 | 0.00518 |
| `zsasa` f32 | 1,024 | 1.000000 | 0.0000964 | 0.00287 |
| RustSASA | 64 | 0.999935 | 0.423 | 2.79 |
| RustSASA | 128 | 0.999973 | 0.275 | 2.15 |
| RustSASA | 256 | 0.999992 | 0.154 | 1.03 |
| RustSASA | 512 | 0.999997 | 0.0896 | 0.682 |
| RustSASA | 1,024 | 0.999999 | 0.0549 | 0.361 |
| Lahuta | 64 | 0.999935 | 0.423 | 2.79 |
| Lahuta | 128 | 0.999973 | 0.275 | 2.15 |
| Lahuta | 256 | 0.999992 | 0.154 | 1.03 |
| Lahuta | 512 | 0.999997 | 0.0897 | 0.684 |
| Lahuta | 1,024 | 0.999999 | 0.0549 | 0.360 |

**Table S3. Static validation, bitmask modes.** Corrected rows use the experimental bias-correction option of `zsasa` v0.12.1. Lahuta's bitmask mode supports 64, 128, and 256 points.

| Tool / mode | Points | Residual $R^2$ | Mean rel. diff. (%) | Max rel. diff. (%) |
|---|---|---|---|---|
| `zsasa` bitmask f64 | 64 | 0.999755 | 0.758 | 2.41 |
| `zsasa` bitmask f64 | 128 | 0.999811 | 0.662 | 2.02 |
| `zsasa` bitmask f64 | 256 | 0.999789 | 0.719 | 1.66 |
| `zsasa` bitmask f64 | 512 | 0.999784 | 0.740 | 1.31 |
| `zsasa` bitmask f64 | 1,024 | 0.999788 | 0.732 | 1.08 |
| `zsasa` bitmask f32 | 64 | 0.999755 | 0.758 | 2.41 |
| `zsasa` bitmask f32 | 128 | 0.999811 | 0.662 | 2.02 |
| `zsasa` bitmask f32 | 256 | 0.999789 | 0.719 | 1.66 |
| `zsasa` bitmask f32 | 512 | 0.999784 | 0.740 | 1.31 |
| `zsasa` bitmask f32 | 1,024 | 0.999788 | 0.732 | 1.08 |
| `zsasa` bitmask f64, corrected | 64 | 0.999748 | 0.777 | 2.42 |
| `zsasa` bitmask f64, corrected | 128 | 0.999716 | 0.858 | 2.23 |
| `zsasa` bitmask f64, corrected | 256 | 0.999763 | 0.794 | 1.63 |
| `zsasa` bitmask f64, corrected | 512 | 0.999777 | 0.772 | 1.49 |
| `zsasa` bitmask f64, corrected | 1,024 | 0.999776 | 0.780 | 1.18 |
| `zsasa` bitmask f32, corrected | 64 | 0.999747 | 0.777 | 2.45 |
| `zsasa` bitmask f32, corrected | 128 | 0.999716 | 0.858 | 2.25 |
| `zsasa` bitmask f32, corrected | 256 | 0.999763 | 0.794 | 1.63 |
| `zsasa` bitmask f32, corrected | 512 | 0.999777 | 0.772 | 1.49 |
| `zsasa` bitmask f32, corrected | 1,024 | 0.999776 | 0.780 | 1.18 |
| Lahuta bitmask | 64 | 0.999701 | 0.841 | 3.04 |
| Lahuta bitmask | 128 | 0.999768 | 0.732 | 2.47 |
| Lahuta bitmask | 256 | 0.999777 | 0.737 | 1.77 |

**Table S4. Static validation, Lee–Richards 20-slice check.** `zsasa` v0.12.1 against the pinned FreeSASA 20-slice output. LR, Lee–Richards.

| Tool / mode | Slices | Shared structures | Residual $R^2$ | Mean signed diff. (Å²) | Mean rel. diff. (%) | Max rel. diff. (%) |
|---|---|---|---|---|---|---|
| `zsasa` LR f64 | 20 | 4,370 | 1.000000 | 0.0 | 0.0000210 | 0.000231 |
| `zsasa` LR f32 | 20 | 4,370 | 1.000000 | 0.0 | 0.0000223 | 0.000229 |

**Table S5. Batch 10-thread results, *E. coli*.** Aggregate timing and memory readouts for the 4,370-structure *E. coli* AlphaFold Database collection.

| Tool / mode | Runtime (s) | Structures/s | ms/structure | Peak RSS (MiB) | Structures/s per MiB |
|---|---|---|---|---|---|
| `zsasa` f64 | 4.457 | 980.6 | 1.020 | 54.2 | 18.10 |
| `zsasa` f32 | 4.380 | 997.7 | 1.002 | 52.1 | 19.16 |
| `zsasa` bitmask f64 | 1.569 | 2,784 | 0.3591 | 56.2 | 49.58 |
| `zsasa` bitmask f32 | 1.526 | 2,864 | 0.3492 | 54.8 | 52.21 |
| FreeSASA batch | 13.85 | 315.6 | 3.168 | 228.8 | 1.380 |
| RustSASA | 6.145 | 711.2 | 1.406 | 170.2 | 4.178 |
| Lahuta | 7.039 | 620.8 | 1.611 | 187.5 | 3.312 |
| Lahuta bitmask | 2.071 | 2,110 | 0.4740 | 176.8 | 11.93 |

**Table S6. Batch thread scaling, *E. coli*.** Speedup relative to one thread and parallel efficiency on the 4,370-structure *E. coli* AlphaFold Database collection; the test machine has 4 performance and 6 efficiency cores. The runs with 20 and 40 `zsasa` workers and the full per-thread readouts are in the benchmark archive.

| Tool / mode | Runtime, 1 thread (s) | Speedup, 4 / 8 / 10 threads | Parallel efficiency, 4 / 8 / 10 threads | Peak RSS, 1 / 10 threads (MiB) |
|---|---|---|---|---|
| `zsasa` f64 | 29.41 | 3.79 / 5.82 / 6.60 | 0.95 / 0.73 / 0.66 | 12.5 / 54.2 |
| `zsasa` f32 | 28.88 | 3.77 / 5.84 / 6.59 | 0.94 / 0.73 / 0.66 | 12.1 / 52.1 |
| `zsasa` bitmask f64 | 10.25 | 3.90 / 5.90 / 6.53 | 0.97 / 0.74 / 0.65 | 14.8 / 56.2 |
| `zsasa` bitmask f32 | 10.22 | 3.91 / 6.02 / 6.70 | 0.98 / 0.75 / 0.67 | 14.4 / 54.8 |
| FreeSASA batch | 42.82 | 3.50 / 3.09 / 3.09 | 0.88 / 0.39 / 0.31 | 86.2 / 228.8 |
| RustSASA | 31.73 | 3.97 / 5.43 / 5.16 | 0.99 / 0.68 / 0.52 | 39.0 / 170.2 |
| Lahuta | 47.24 | 3.90 / 5.95 / 6.71 | 0.97 / 0.74 / 0.67 | 84.1 / 187.5 |
| Lahuta bitmask | 13.42 | 4.01 / 5.92 / 6.48 | 1.00 / 0.74 / 0.65 | 82.1 / 176.8 |

**Table S7. Worker counts above the CPU count, Human and Swiss-Prot collections.** Batch runs with `zsasa` v0.12.1 at 128 sphere points on a machine with 10 logical CPUs. Human rows are means of three runs. Swiss-Prot rows (550,122 structures) were measured once without warmup under the passive disk I/O policy of macOS; the rows marked default I/O policy repeat them, together with Lahuta, in a separate session (Supplementary Note S2). Comparators were measured at 10 threads only. Busy cores is the CPU proxy of Table S1.

| Collection | Tool / mode | Workers | Runtime (s) | Structures/s | Peak RSS (MiB) | Busy cores |
|---|---|---|---|---|---|---|
| Human | `zsasa` f64 | 10 | 44.20 | 533.6 | 96.2 | 9.4 |
| Human | `zsasa` f64 | 20 | 43.96 | 536.5 | 178.3 | 9.6 |
| Human | `zsasa` f64 | 40 | 43.90 | 537.3 | 323.8 | 9.6 |
| Human | `zsasa` f32 | 10 | 43.44 | 543.0 | 91.3 | 9.4 |
| Human | `zsasa` f32 | 20 | 44.43 | 530.9 | 166.8 | 9.2 |
| Human | `zsasa` f32 | 40 | 42.97 | 548.9 | 311.6 | 9.6 |
| Human | `zsasa` bitmask f64 | 10 | 14.73 | 1,601 | 98.5 | 8.8 |
| Human | `zsasa` bitmask f64 | 20 | 14.33 | 1,646 | 177.8 | 9.3 |
| Human | `zsasa` bitmask f64 | 40 | 14.39 | 1,639 | 327.2 | 9.2 |
| Human | `zsasa` bitmask f32 | 10 | 14.35 | 1,644 | 93.6 | 8.8 |
| Human | `zsasa` bitmask f32 | 20 | 13.96 | 1,690 | 169.2 | 9.3 |
| Human | `zsasa` bitmask f32 | 40 | 14.04 | 1,679 | 313.7 | 9.2 |
| Human | FreeSASA batch | 10 | 147.2 | 160.2 | 676.4 | 7.9 |
| Human | RustSASA | 10 | 56.21 | 419.6 | 328.5 | 7.2 |
| Human | Lahuta | 10 | 71.55 | 329.6 | 340.5 | 9.5 |
| Human | Lahuta bitmask | 10 | 19.81 | 1,191 | 352.5 | 9.3 |
| Swiss-Prot | `zsasa` f32 | 10 | 748.8 | 734.7 | 289.2 | 7.5 |
| Swiss-Prot | `zsasa` f32 | 20 | 708.8 | 776.2 | 370.6 | 8.2 |
| Swiss-Prot | `zsasa` f32 | 40 | 710.2 | 774.6 | 527.4 | 8.1 |
| Swiss-Prot | `zsasa` bitmask f32 | 10 | 344.0 | 1,599 | 291.3 | 4.8 |
| Swiss-Prot | `zsasa` bitmask f32 | 20 | 330.3 | 1,666 | 372.8 | 5.2 |
| Swiss-Prot | `zsasa` bitmask f32 | 40 | 351.3 | 1,566 | 530.8 | 4.9 |
| Swiss-Prot | Lahuta bitmask | 10 | 433.5 | 1,269 | 659.5 | 5.7 |
| Swiss-Prot, default I/O policy | `zsasa` f32 | 10 | 864.8 | 636.2 | 289.4 | 5.7 |
| Swiss-Prot, default I/O policy | `zsasa` f32 | 20 | 713.9 | 770.6 | 369.0 | 7.9 |
| Swiss-Prot, default I/O policy | `zsasa` f32 | 40 | 701.6 | 784.1 | 529.9 | 8.1 |

| Collection | Tool / mode | Workers | Runtime (s) | Structures/s | Peak RSS (MiB) | Busy cores |
|---|---|---|---|---|---|---|
| Swiss-Prot, default I/O policy | zsasa bitmask f32 | 10 | 534.5 | 1,029 | 291.5 | 2.9 |
| Swiss-Prot, default I/O policy | zsasa bitmask f32 | 20 | 405.8 | 1,356 | 372.7 | 4.3 |
| Swiss-Prot, default I/O policy | zsasa bitmask f32 | 40 | 377.1 | 1,459 | 531.9 | 4.8 |
| Swiss-Prot, default I/O policy | Lahuta bitmask | 10 | 600.7 | 915.7 | 673.3 | 3.7 |

**Table S8. Batch 10-thread results, Human.** Aggregate timing and memory readouts for the 23,586-structure human AlphaFold Database collection.

| Tool / mode | Runtime (s) | Structures/s | ms/structure | Peak RSS (MiB) | Structures/s per MiB |
|---|---|---|---|---|---|
| zsasa f64 | 44.20 | 533.6 | 1.874 | 96.2 | 5.549 |
| zsasa f32 | 43.44 | 543.0 | 1.842 | 91.3 | 5.947 |
| zsasa bitmask f64 | 14.73 | 1,601 | 0.6245 | 98.5 | 16.26 |
| zsasa bitmask f32 | 14.35 | 1,644 | 0.6084 | 93.6 | 17.56 |
| FreeSASA batch | 147.2 | 160.2 | 6.243 | 676.4 | 0.2368 |
| RustSASA | 56.21 | 419.6 | 2.383 | 328.5 | 1.277 |
| Lahuta | 71.55 | 329.6 | 3.034 | 340.5 | 0.9682 |
| Lahuta bitmask | 19.81 | 1,191 | 0.8397 | 352.5 | 3.379 |

**Table S9. Single-file subset.**

| Structure | Source | Atoms | Chains | Role |
|---|---|---|---|---|
| AF-P49792-F10 | AFDB single chain | 10,919 | 1 | Medium single-chain case |
| AF-Q6ZS30-F1 | AFDB single chain | 21,611 | 1 | Large single-chain case |
| AF-0000000066638622 | AFDB-derived complex | 14,618 | 2 | Medium two-chain case |
| AF-0000000065781219 | AFDB-derived complex | 24,140 | 2 | Large two-chain case |
| 3jc8 | PDB assembly | 107,500 | 115 | 100,000-atom case |
| 5vyc | PDB assembly | 249,168 | 216 | Large multi-chain case |
| 8rbs | PDB assembly | 164,605 | 85 | Coordinates above 999.999 Å |
| 9fqr | PDB assembly | 4,506,416 | 1,574 | Maximum-size case; coordinates above 1,000 Å |

**Table S10. Single-file 10-thread results.** Aggregate timing and memory readouts for the curated single-file subset (mmCIF input, 100 sphere points), one block per structure. PDBTools.jl runtime includes Julia startup. FreeSASA stopped without assigning atomic radii on the two AFDB-derived complexes, so those rows are empty.

**AF-P49792-F10** (10,919 atoms, 1 chain)

| Tool / mode | Runtime (s) | Atoms/s | Peak RSS (MiB) |
|---|---|---|---|
| zsasa f64 | 0.02218 | 492,324 | 8.0 |
| zsasa f32 | 0.02200 | 496,311 | 8.0 |
| zsasa bitmask f64 | 0.03394 | 321,723 | 9.1 |
| zsasa bitmask f32 | 0.03395 | 321,648 | 8.6 |
| FreeSASA | 0.05406 | 201,985 | 66.6 |
| RustSASA | 0.07808 | 139,845 | 29.8 |
| PDBTools.jl | 1.906 | 5,728 | 466.9 |

**AF-0000000066638622** (14,618 atoms, 2 chains)

| Tool / mode | Runtime (s) | Atoms/s | Peak RSS (MiB) |
|---|---|---|---|
| zsasa f64 | 0.03110 | 469,979 | 9.8 |
| zsasa f32 | 0.03095 | 472,346 | 9.8 |
| zsasa bitmask f64 | 0.04076 | 358,634 | 10.3 |
| zsasa bitmask f32 | 0.04077 | 358,567 | 10.4 |
| FreeSASA | – | – | – |
| RustSASA | 0.04426 | 330,294 | 30.5 |
| PDBTools.jl | 1.748 | 8,362 | 455.4 |

**AF-Q6ZS30-F1** (21,611 atoms, 1 chain)

| Tool / mode | Runtime (s) | Atoms/s | Peak RSS (MiB) |
|---|---|---|---|
| zsasa f64 | 0.03907 | 553,109 | 9.4 |
| zsasa f32 | 0.03906 | 553,226 | 9.4 |
| zsasa bitmask f64 | 0.04950 | 436,578 | 12.5 |
| zsasa bitmask f32 | 0.04896 | 441,392 | 12.1 |
| FreeSASA | 0.1114 | 193,981 | 130.0 |
| RustSASA | 0.08681 | 248,950 | 66.9 |
| PDBTools.jl | 1.917 | 11,275 | 479.7 |

**AF-0000000065781219** (24,140 atoms, 2 chains)

| Tool / mode | Runtime (s) | Atoms/s | Peak RSS (MiB) |
|---|---|---|---|
| zsasa f64 | 0.04110 | 587,375 | 10.4 |
| zsasa f32 | 0.04256 | 567,163 | 10.1 |
| zsasa bitmask f64 | 0.04991 | 483,694 | 13.6 |
| zsasa bitmask f32 | 0.05160 | 467,811 | 13.1 |
| FreeSASA | – | – | – |
| RustSASA | 0.07316 | 329,964 | 35.1 |
| PDBTools.jl | 1.787 | 13,507 | 467.4 |

**3jc8** (107,500 atoms, 115 chains)

| Tool / mode | Runtime (s) | Atoms/s | Peak RSS (MiB) |
|---|---|---|---|
| zsasa f64 | 0.1459 | 736,605 | 40.9 |
| zsasa f32 | 0.1469 | 732,012 | 41.1 |
| zsasa bitmask f64 | 0.1404 | 765,581 | 42.9 |
| zsasa bitmask f32 | 0.1391 | 772,998 | 41.1 |
| FreeSASA | 0.6414 | 167,597 | 637.6 |
| RustSASA | 0.3249 | 330,854 | 246.8 |
| PDBTools.jl | 1.614 | 66,598 | 619.5 |

**8rbs** (164,605 atoms, 85 chains)

| Tool / mode | Runtime (s) | Atoms/s | Peak RSS (MiB) |
|---|---|---|---|
| zsasa f64 | 0.2094 | 785,970 | 57.0 |
| zsasa f32 | 0.2086 | 789,143 | 55.9 |
| zsasa bitmask f64 | 0.1948 | 844,819 | 59.2 |
| zsasa bitmask f32 | 0.1934 | 850,945 | 55.9 |
| FreeSASA | 0.9333 | 176,363 | 966.5 |
| RustSASA | 0.4808 | 342,334 | 315.4 |
| PDBTools.jl | 1.725 | 95,441 | 676.0 |

**5vyc** (249,168 atoms, 216 chains)

| Tool / mode | Runtime (s) | Atoms/s | Peak RSS (MiB) |
|---|---|---|---|
| zsasa f64 | 0.3830 | 650,579 | 124.3 |
| zsasa f32 | 0.3823 | 651,779 | 124.1 |
| zsasa bitmask f64 | 0.3472 | 717,601 | 126.2 |
| zsasa bitmask f32 | 0.3448 | 722,743 | 124.2 |
| FreeSASA | 2.058 | 121,070 | 1,822.3 |
| RustSASA | 1.070 | 232,819 | 734.6 |
| PDBTools.jl | 1.892 | 131,703 | 776.2 |

**9fqr** (4,506,416 atoms, 1,574 chains)

**Table S11. Comparator-specific caveats.**

| Tool / mode | Runtime (s) | Atoms/s | Peak RSS (MiB) |
|---|---|---|---|
| zsasa f64 | 6.266 | 719,166 | 2,191.7 |
| zsasa f32 | 6.356 | 709,038 | 2,201.1 |
| zsasa bitmask f64 | 5.357 | 841,236 | 2,228.2 |
| zsasa bitmask f32 | 5.324 | 846,424 | 2,203.2 |
| FreeSASA | 28.95 | 155,656 | 20,871.7 |
| RustSASA | 23.65 | 190,571 | 8,212.0 |
| PDBTools.jl | 10.51 | 428,832 | 6,728.8 |

**Table S12. Molecular-dynamics validation, command-line bitmask modes.** Pearson $r^2$ remains high, but the uncorrected bitmask totals are systematically lower than the MDTraj reference, so residual $R^2$ is much smaller. Corrected rows use the experimental bias-correction option. Rows use the single-lookup-table mode.

| Comparator or context | Caveat |
|---|---|
| FreeSASA batch | Uses a wrapper around the FreeSASA C API |
| FreeSASA single file | Native single-file command |
| RustSASA totals | Protein-level JSON output used |
| FreeSASA mmCIF input | Invoked with `--cif`; stale assembly-generation metadata that the pinned parser cannot read was removed |
| Lahuta batch | Included for AFDB-style batch inputs |
| Lahuta single file | Excluded from the mostly multi-chain subset |
| MDTraj validation | MDTraj uses the same golden-section spiral of sphere points as `zsasa` and FreeSASA but with the $y$ axis as the polar axis; `zsasa` and FreeSASA use the $z$ axis |
| RustSASA validation | RustSASA uses a golden-section spiral with the $z$ axis as the polar axis, shifted by half a step relative to FreeSASA and wound in the opposite direction |
| MDTraj reference | Computed one frame at a time; a multi-frame call to MDTraj's Shrake–Rupley function inflates the areas of every frame after the first (by 1.5% at 64 points and 0.09% at 1,024 points on 5wvo_C; reported at https://github.com/mdtraj/mdtraj/issues/2195) |
| MDTraj validation, radii | MDTraj and the `zsasa` + MDTraj integration use MDTraj's element radii; the `zsasa` CLI uses NACCESS radii and the `zsasa` + MDAnalysis integration uses MDAnalysis radii |
| mdsasa-bolt memory | Includes wrapper integration behavior |
| Output granularity | Tools emit different detail levels |
| PDBTools.jl runtime | Full-process runtime includes Julia startup; component timings from a warm process |
| Comparator timings | Validation, batch, and trajectory comparators measured in the `zsasa` v0.12.1 session; single-file comparator timings and mdsasa-bolt on 5vz0_A retained from earlier sessions on the same machine (FreeSASA and RustSASA on PDB input from the `zsasa` v0.6.0 session, all other rows from the v0.9.0 session) |

**Table S13. Molecular-dynamics validation, exact modes and Python integrations.** Per-frame total SASA against MDTraj on the 1,001 frames of 5wvo_C. Residual $R^2$ measures absolute agreement and penalizes systematic offsets; Pearson $r^2$ measures frame-to-frame co-variation.

| Tool / path | Points | Residual $R^2$ | Pearson $r^2$ | Mean signed diff. (Å$^2$) | Mean rel. diff. (%) | Max rel. diff. (%) |
|---|---|---|---|---|---|---|
| `zsasa` CLI bitmask f64 | 64 | 0.459397 | 0.964406 | −302.4 | 2.02 | 3.72 |
| `zsasa` CLI bitmask f64 | 128 | 0.473926 | 0.985864 | −300.1 | 2.00 | 2.93 |
| `zsasa` CLI bitmask f64 | 256 | 0.457150 | 0.992350 | −306.0 | 2.04 | 2.74 |
| `zsasa` CLI bitmask f64 | 512 | 0.459839 | 0.995791 | −306.2 | 2.05 | 2.68 |
| `zsasa` CLI bitmask f64 | 1,024 | 0.464143 | 0.996703 | −305.3 | 2.04 | 2.58 |
| `zsasa` CLI bitmask f32 | 64 | 0.459409 | 0.964381 | −302.4 | 2.02 | 3.73 |
| `zsasa` CLI bitmask f32 | 128 | 0.473927 | 0.985871 | −300.1 | 2.00 | 2.94 |
| `zsasa` CLI bitmask f32 | 256 | 0.457244 | 0.992344 | −306.0 | 2.04 | 2.74 |
| `zsasa` CLI bitmask f32 | 512 | 0.459935 | 0.995797 | −306.2 | 2.05 | 2.68 |
| `zsasa` CLI bitmask f32 | 1,024 | 0.464191 | 0.996707 | −305.3 | 2.04 | 2.58 |
| `zsasa` CLI bitmask f64, corrected | 64 | 0.946056 | 0.964399 | −57.6 | 0.536 | 2.12 |
| `zsasa` CLI bitmask f64, corrected | 128 | 0.968707 | 0.985866 | −54.7 | 0.413 | 1.30 |
| `zsasa` CLI bitmask f64, corrected | 256 | 0.971160 | 0.992327 | −60.7 | 0.417 | 1.11 |
| `zsasa` CLI bitmask f64, corrected | 512 | 0.974478 | 0.995770 | −60.8 | 0.408 | 1.05 |

| Tool / path | Points | Residual $R^2$ | Pearson $r^2$ | Mean signed diff. (Å$^2$) | Mean rel. diff. (%) | Max rel. diff. (%) |
|---|---|---|---|---|---|---|
| zsasa CLI bitmask f64, corrected | 1,024 | 0.976042 | 0.996682 | −59.9 | 0.400 | 0.950 |
| zsasa CLI bitmask f32, corrected | 64 | 0.946038 | 0.964374 | −57.6 | 0.536 | 2.13 |
| zsasa CLI bitmask f32, corrected | 128 | 0.968712 | 0.985872 | −54.7 | 0.413 | 1.31 |
| zsasa CLI bitmask f32, corrected | 256 | 0.971174 | 0.992322 | −60.6 | 0.417 | 1.11 |
| zsasa CLI bitmask f32, corrected | 512 | 0.974501 | 0.995775 | −60.8 | 0.408 | 1.05 |
| zsasa CLI bitmask f32, corrected | 1,024 | 0.976055 | 0.996686 | −59.9 | 0.400 | 0.949 |

**Table S14. Molecular-dynamics throughput results.** Aggregate trajectory-level timing and memory readouts for the three MD workloads at 128 sphere points and 10 threads (zsasa v0.12.1). Command-line bitmask rows use the single-lookup-table mode without bias correction.

| Tool / path | Points | Residual $R^2$ | Pearson $r^2$ | Mean signed diff. (Å$^2$) | Mean rel. diff. (%) | Max rel. diff. (%) |
|---|---|---|---|---|---|---|
| zsasa + MDTraj | 64 | 0.972891 | 0.973069 | 4.6 | 0.376 | 1.62 |
| zsasa + MDTraj | 128 | 0.989363 | 0.989383 | 0.4 | 0.230 | 1.29 |
| zsasa + MDTraj | 256 | 0.995994 | 0.995994 | −0.1 | 0.142 | 0.686 |
| zsasa + MDTraj | 512 | 0.998609 | 0.998609 | 0.1 | 0.0832 | 0.366 |
| zsasa + MDTraj | 1,024 | 0.999515 | 0.999516 | −0.2 | 0.0492 | 0.241 |
| zsasa + MDAnalysis | 64 | 0.909662 | 0.967514 | 102.3 | 0.725 | 2.18 |
| zsasa + MDAnalysis | 128 | 0.935340 | 0.987186 | 95.6 | 0.643 | 1.88 |
| zsasa + MDAnalysis | 256 | 0.940582 | 0.993927 | 96.7 | 0.647 | 1.45 |
| zsasa + MDAnalysis | 512 | 0.944244 | 0.997055 | 96.2 | 0.644 | 1.09 |
| zsasa + MDAnalysis | 1,024 | 0.945322 | 0.997766 | 95.9 | 0.642 | 1.05 |
| zsasa CLI f64 | 64 | 0.964029 | 0.967036 | −22.3 | 0.436 | 1.75 |
| zsasa CLI f64 | 128 | 0.981550 | 0.986503 | −29.3 | 0.303 | 1.12 |
| zsasa CLI f64 | 256 | 0.988984 | 0.993638 | −28.4 | 0.242 | 0.928 |
| zsasa CLI f64 | 512 | 0.991685 | 0.996278 | −27.7 | 0.212 | 0.709 |
| zsasa CLI f64 | 1,024 | 0.992473 | 0.997159 | −28.0 | 0.204 | 0.656 |
| zsasa CLI f32 | 64 | 0.964047 | 0.967048 | −22.3 | 0.436 | 1.75 |
| zsasa CLI f32 | 128 | 0.981550 | 0.986501 | −29.3 | 0.303 | 1.12 |
| zsasa CLI f32 | 256 | 0.988985 | 0.993638 | −28.4 | 0.242 | 0.928 |
| zsasa CLI f32 | 512 | 0.991685 | 0.996277 | −27.7 | 0.212 | 0.709 |
| zsasa CLI f32 | 1,024 | 0.992476 | 0.997160 | −28.0 | 0.204 | 0.656 |

**Table S15. AFDB dimer workflow runs and ΔSASA summary.** Both runs used `zsasa` v0.12.1 in bitmask f32 mode at 128 sphere points, with the CCD classifier, a 1.4 Å probe, and 20 workers on the Apple M4 laptop (10 cores, 32 GB) used for the benchmarks. Each set was run once under the default disk I/O policy of macOS. The rows marked passive policy repeat each run under the passive disk I/O policy (Supplementary Note S2); the repeated runs gave the same results.

| Dataset | Frames | Atoms | Tool / mode | Runtime (s) | Frames/s | Peak RSS (MiB) | Frames/s per MiB |
|---|---|---|---|---|---|---|---|
| 5wvo_C | 1,001 | 3,858 | zsasa CLI f64 | 2.098 | 477.2 | 25.5 | 18.74 |
| 5wvo_C | 1,001 | 3,858 | zsasa CLI f32 | 2.122 | 471.8 | 25.3 | 18.68 |
| 5wvo_C | 1,001 | 3,858 | zsasa CLI bitmask f64 | 0.8891 | 1,126 | 25.4 | 44.28 |
| 5wvo_C | 1,001 | 3,858 | zsasa CLI bitmask f32 | 0.8529 | 1,174 | 25.5 | 46.08 |
| 5wvo_C | 1,001 | 3,858 | zsasa + MDTraj | 2.074 | 482.7 | 210.4 | 2.294 |
| 5wvo_C | 1,001 | 3,858 | zsasa + MDTraj bitmask | 0.8803 | 1,137 | 209.3 | 5.433 |
| 5wvo_C | 1,001 | 3,858 | zsasa + MDAnalysis | 2.301 | 435.1 | 193.2 | 2.251 |
| 5wvo_C | 1,001 | 3,858 | zsasa + MDAnalysis bitmask | 1.146 | 873.3 | 192.7 | 4.533 |
| 5wvo_C | 1,001 | 3,858 | MDTraj | 26.22 | 38.17 | 160.2 | 0.2383 |
| 5wvo_C | 1,001 | 3,858 | mdsasa-bolt (Rust) | 4.475 | 223.7 | 1,416.1 | 0.1579 |
| 6sup_A | 1,001 | 33,377 | zsasa CLI f64 | 18.10 | 55.31 | 153.2 | 0.3609 |
| 6sup_A | 1,001 | 33,377 | zsasa CLI f32 | 17.29 | 57.88 | 136.0 | 0.4256 |
| 6sup_A | 1,001 | 33,377 | zsasa CLI bitmask f64 | 7.132 | 140.4 | 152.4 | 0.9210 |
| 6sup_A | 1,001 | 33,377 | zsasa CLI bitmask f32 | 6.945 | 144.1 | 142.8 | 1.009 |
| 6sup_A | 1,001 | 33,377 | zsasa + MDTraj | 18.07 | 55.38 | 1,353.2 | 0.04093 |
| 6sup_A | 1,001 | 33,377 | zsasa + MDTraj bitmask | 6.974 | 143.5 | 1,343.5 | 0.1068 |
| 6sup_A | 1,001 | 33,377 | zsasa + MDAnalysis | 18.12 | 55.25 | 960.1 | 0.05755 |
| 6sup_A | 1,001 | 33,377 | zsasa + MDAnalysis bitmask | 7.330 | 136.6 | 956.6 | 0.1428 |
| 6sup_A | 1,001 | 33,377 | MDTraj | 944.9 | 1.059 | 1,002.8 | 0.001056 |
| 6sup_A | 1,001 | 33,377 | mdsasa-bolt (Rust) | 59.30 | 16.88 | 11,658.0 | 0.001448 |
| 5vz0_A | 10,001 | 17,910 | zsasa CLI f64 | 96.86 | 103.3 | 87.1 | 1.186 |
| 5vz0_A | 10,001 | 17,910 | zsasa CLI f32 | 94.07 | 106.3 | 81.7 | 1.301 |
| 5vz0_A | 10,001 | 17,910 | zsasa CLI bitmask f64 | 38.41 | 260.4 | 88.0 | 2.960 |
| 5vz0_A | 10,001 | 17,910 | zsasa CLI bitmask f32 | 37.67 | 265.5 | 82.3 | 3.227 |
| 5vz0_A | 10,001 | 17,910 | mdsasa-bolt (Rust) | 2,508 | 3.988 | 24,332.9 | 0.0001639 |

**Table S16. Software and comparator versions.** Versions were pinned before the benchmark evidence was collected.

| Quantity | Human homodimers | Human–human heterodimers |
|---|---|---|
| Models processed | 91,637 | 531,605 |
| Failed models | 0 | 0 |
| Wall time (s) | 177.1 | 1,418.8 |
| User CPU time (s) | 1,483.6 | 12,315.7 |
| Peak RSS (MiB) | 1,655.0 | 350.1 |
| Wall time, passive policy (s) | 177.4 | 1,404.6 |
| Peak RSS, passive policy (MiB) | 1,668.4 | 345.4 |
| Median ΔSASA (Å²) | 2,829 | 2,933 |
| Interquartile range of ΔSASA (Å²) | 1,807–4,536 | 1,793–4,641 |
| 95th percentile of ΔSASA (Å²) | 9,882 | 8,433 |
| Models with ΔSASA = 0 | 879 (1.0%) | 9,288 (1.7%) |
| Models with ΔSASA < 0 | 0 | 0 |
| High-quality models | 5.1% | 0.8% |

| Quantity | Human homodimers | Human–human heterodimers |
|---|---|---|
| Models with $ipSAE_{min} = 0$ | 53.1% | 79.6% |
| Spearman $\rho$, $\Delta$SASA versus mean pLDDT | −0.01 | −0.32 |
| Spearman $\rho$, $\Delta$SASA versus $ipSAE_{min}$ | 0.24 | 0.25 |
| Spearman $\rho$, $\Delta$SASA versus ipTM | 0.37 | 0.43 |
| Spearman $\rho$, $\Delta$SASA versus pDockQ | 0.46 | 0.53 |

**Table S17. Benchmark inputs and timing protocol.** A filesystem `sync` preceded every timed run. Validation runs were not timed.

| Tool | Version | Role in this study |
|---|---|---|
| `zsasa` | v0.12.1 | Method under evaluation |
| FreeSASA | 2.1.3 | Static-validation reference; batch comparator through wrapper |
| RustSASA | 0.9.2 | Batch and single-file comparator |
| Lahuta | 2.0.3 | Batch comparator |
| MDTraj | 1.11.1 | Trajectory-validation reference and trajectory comparator |
| MDAnalysis | 2.10.0 | Trajectory integration dependency |
| mdsasa-bolt (Campbell 2026) | 4.0.1 | Trajectory comparator; MDAnalysis interface to RustSASA (https://github.com/maxall41/mdsasa-bolt) |
| PDBTools.jl (Martínez and Lima 2026) | 3.33.2 | Single-file comparator |

**Table S18. Modes, comparators, and settings.** Trajectory suites used stride 1 with hydrogens included.

| Workload | Input | Sphere points | Threads | Warmup / measured runs |
|---|---|---|---|---|
| Static validation | AFDB v6 *E. coli* K-12 proteome, PDB; 4,370 structures | 64, 128, 256, 512, 1,024; Lee–Richards 20 slices | 10 | Not timed |
| Batch, *E. coli* | AFDB v6 *E. coli* K-12 proteome, PDB; 4,370 structures | 128 | 1, 4, 8, 10; `zsasa` also 20 and 40 workers | 3 / 3 |
| Batch, Human | AFDB v6 human proteome, PDB; 23,586 structures | 128 | 10; `zsasa` also 20 and 40 workers | 3 / 3 |
| Batch, Swiss-Prot | AFDB v6 Swiss-Prot, PDB; 550,122 structures | 128 | `zsasa` 10, 20, and 40 workers; Lahuta bitmask 10 | 0 / 1 |
| Single file | 8 AFDB and PDB structures, uncompressed mmCIF (Table S9) | 100 | 1, 4, 8, 10 | 1 / 3 |
| Trajectory validation | ATLAS 5wvo_C; 3,858 atoms, 1,001 frames | 64, 128, 256, 512, 1,024 | 10 where applicable | Not timed |
| Trajectory throughput | ATLAS 5wvo_C (3,858 atoms, 1,001 frames), 6sup_A (33,377 atoms, 1,001 frames), and 5vz0_A (17,910 atoms, 10,001 frames) | 128 | 10 | 1 / 3 |

**Table S19. Benchmark execution environment.** Head-to-head benchmarks were run on the M4 system unless a row explicitly states otherwise.

| Suite | `zsasa` modes | Comparators | Settings |
|---|---|---|---|
| Static validation | f64, f32; bitmask f64 and f32, each with and without the experimental bias correction; Lee–Richards f64 and f32 | FreeSASA (reference), RustSASA, Lahuta, Lahuta bitmask | Total SASA per structure |
| Batch | f64, f32, bitmask f64, bitmask f32 | FreeSASA batch wrapper, RustSASA, Lahuta, Lahuta bitmask; Swiss-Prot: Lahuta bitmask only | Swiss-Prot under the passive disk I/O policy of macOS, repeated under the default policy (Supplementary Note S2) |
| Single file | f64, f32, bitmask f64, bitmask f32 | FreeSASA, RustSASA, PDBTools.jl; Lahuta excluded | Wall-clock and component (parse and SASA) timing |
| Trajectory validation | CLI exact and bitmask modes; Python MDTraj and MDAnalysis integrations | MDTraj (reference) | NACCESS radii for the CLI; total SASA per frame |
| Trajectory throughput | CLI f64, f32, bitmask f64, bitmask f32 (single lookup table, no bias correction); Python MDTraj and MDAnalysis integrations | Native MDTraj, mdsasa-bolt; 5vz0_A: `zsasa` CLI and mdsasa-bolt only | NACCESS radii for the CLI |

| Category | Setting | Value |
|---|---|---|
| Hardware | Computer | MacBook Pro, Mac16,1 |
| Hardware | Processor | Apple M4 |
| Hardware | CPU cores | 10 total, 4 performance and 6 efficiency |
| Hardware | Memory | 32 GB |
| Operating system | Version | macOS 26 |
| Timing | Tool | hyperfine 1.20.0 |
| Timing | Cache preparation | `sync` before timed runs |
| Python | Version | 3.12 |
| Python | Environment manager | uv |
| Native tools | Build environment | Nix |
| Result storage | Database | DuckDB |
| Dimer workflows | Computer | Same M4 system; ordinary runs, not controlled timings |

### Supplementary figures

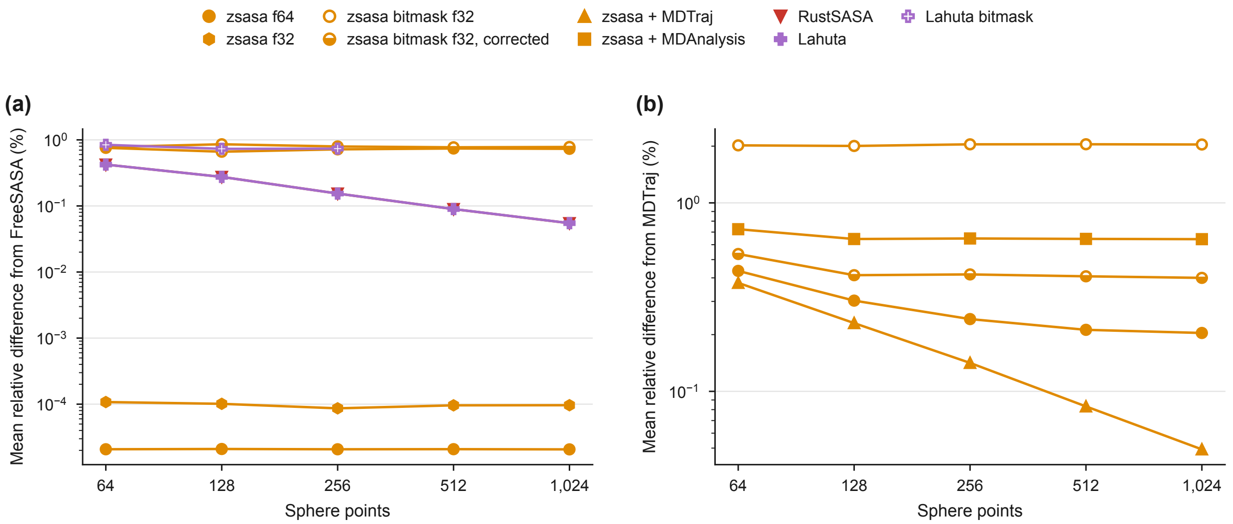


**Figure S1. Validation trends across point counts.** Mean relative difference from the reference as a function of sphere-point count (`zsasa` v0.12.1). (a) Static-structure validation against FreeSASA on 4,370 *E. coli* AlphaFold Database structures. RustSASA and Lahuta nearly coincide, and Lahuta's bitmask mode is shown for its supported 64–256 points. (b) Trajectory validation against MDTraj on the 1,001 frames of 5wvo_C; `zsasa` f64 and bitmask f32 denote the command-line path. Filled markers denote exact modes, open markers bitmask modes, and the half-filled marker the bitmask mode with the experimental bias correction.

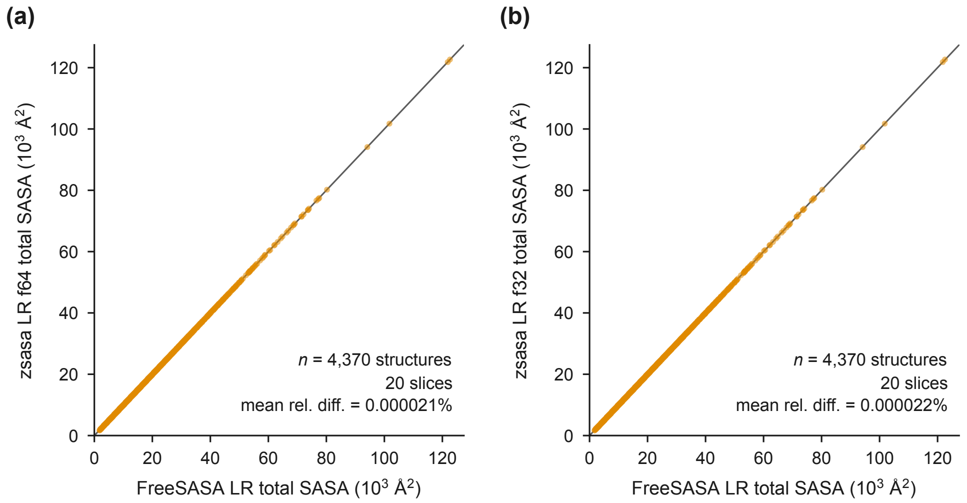


**Figure S2. Lee–Richards validation against FreeSASA.** Per-structure total SASA from 20-slice `zsasa` v0.12.1 Lee–Richards (LR) runs versus 20-slice FreeSASA Lee–Richards output on the 4,370-structure *E. coli* AlphaFold Database validation set. (a) `zsasa` LR f64. (b) `zsasa` LR f32. The line is $y = x$.

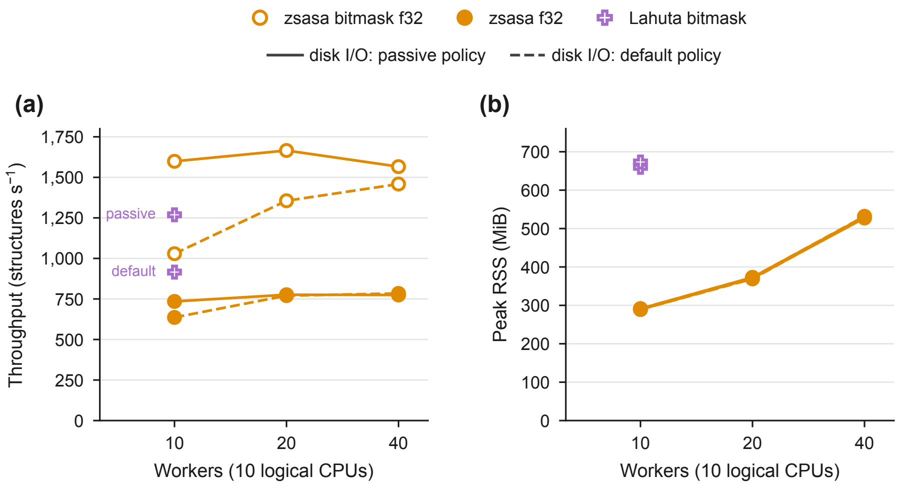


**Figure S3. Worker counts above the CPU count on the Swiss-Prot collection.** (a) Throughput and (b) peak RSS for the 550,122 Swiss-Prot structures of the AlphaFold Database in PDB format at 128 sphere points, with 10, 20, and 40 batch workers on a machine with 10 logical CPUs, under the passive disk I/O policy of macOS (solid lines) and under the default policy (dashed lines; Supplementary Note S2). All `zsasa` runs used v0.12.1. In (b) the curves of the two `zsasa` modes and the two policies lie close together. Lahuta's bitmask mode was measured at 10 threads only, once under each policy. Each condition was measured once, without warmup, so no uncertainty estimate is shown.

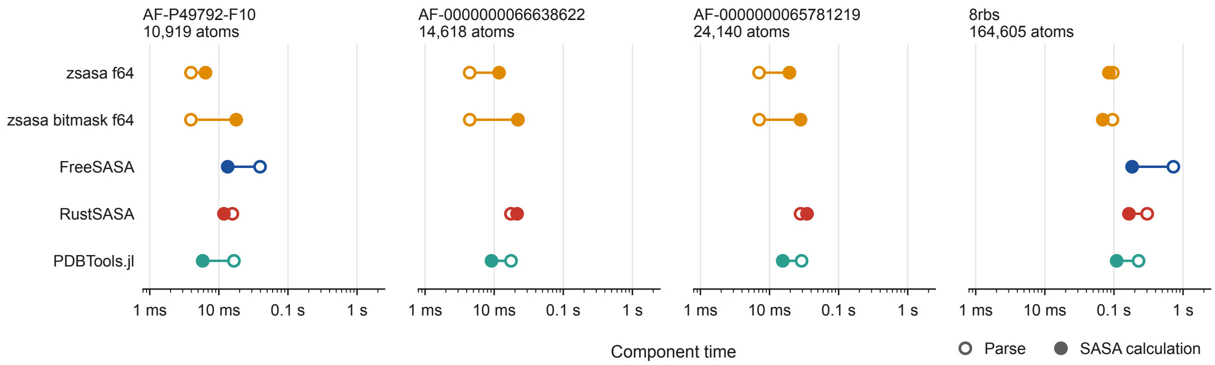


**Figure S4. Additional single-file parse and SASA component timing.** Parse time (open markers) and SASA-calculation time (filled markers) at 10 threads and 100 sphere points for the four single-file structures not shown in Figure 4c of the main text. PDBTools.jl component timings come from a warm process.

Campbell, Maxwell J. 2026. "RustSASA: A Rust Crate for Accelerated Solvent Accessible Surface Area Calculations." *Journal of Open Source Software* 11 (117): 9537. https://doi.org/10.21105/joss.09537.

Lee, B., and F. M. Richards. 1971. "The Interpretation of Protein Structures: Estimation of Static Accessibility." *Journal of Molecular Biology* 55 (3): 379–400. https://doi.org/10.1016/0022-2836(71)90324-X.

Martínez, Leandro, and Ana B B Lima. 2026. "PDBTools.jl: A Lightweight and High-Performance Julia Package for Molecular Structure File Handling and Analysis." *J. Chem. Inf. Model.* 66 (11): 6204–10.

Sejdiu, Besian I., and Madan Babu. 2026. "BPS2026 – High-Performance, Scalable Structural Analysis with Lahuta in the Era of Generative AI." *Biophysical Journal* 125 (4): 233a–234a. https://doi.org/10.1016/j.bpj.2025.11.1538.

Shrake, A., and J. A. Rupley. 1973. "Environment and Exposure to Solvent of Protein Atoms. Lysozyme and Insulin." *Journal of Molecular Biology* 79 (2): 351–71. https://doi.org/10.1016/0022-2836(73)90011-9.
